# Activation dynamics and assembly of root zone soil bacterial communities in response to stress-associated phytohormones

**DOI:** 10.1101/2025.04.23.650272

**Authors:** Sreejata Bandopadhyay, Oishi Bagchi, Ashley Shade

**Affiliations:** The Plant Resilience Institute, Michigan State University, 567 Wilson Rd, East Lansing MI 48824; The Great Lakes Bioenergy Research Center, Michigan State University, 567 Wilson Rd, East Lansing MI 48824; Universite Claude Bernard Lyon 1, CNRS, INRAE, VetAgro Sup, Laboratoire d’Ecologie Microbienne LEM, CNRS UMR5557, INRAE UMR1418, Villeurbanne, F-69100 France

**Keywords:** Plant microbiome, rhizosphere, salicylic acid, abscisic acid, mesocosm, 16S ratios, amplicon sequencing, transcriptionally active populations, reactivation, before after control impact design (BACI)

## Abstract

Plants can “cry for help” to recruit supportive microbiome members during stressful conditions. We evaluated the activation dynamics of root zone soil bacteria in response to phytohormones produced when plants are stressed, hypothesizing that the activated taxa support plant resilience. We conducted a 2-week laboratory experiment using mesocosms of root zone soil collected from two different crops: the annual legume common bean (*Phaseolus vulgaris* L.) and the perennial grass switchgrass (*Panicum virgatum*). We inactivated the microbiome by drying and then treated the soils with either abscisic acid, salicylic acid, a carrier control (methanol), or water, and then quantified the reactivation dynamics of bacterial populations over time, at one, 7, and 14 days after phytohormone addition, using amplicon sequencing of 16S rRNA and rDNA. There were several Actinobacterial taxa that switched from an average population-inactive to a population-active state after exposure to abscisic acid and salicylic acid, with *Microbispora* lineages switching especially noted. Some taxa were activated only in one crop’s soil, and some were activated in both crops’ soils in response to the same phytohormone. This work suggests that different bacteria have different specificities to phytohormones as plant stress signals and provides insights into understanding the mechanisms by which stressed plants may “cry for help” to recruit bacteria from the root zone to the rhizosphere.

**Importance:** Global food security is an urgent societal challenge that has been intensified by climate change and other anthropogenic stressors placed on the environment. Microbial bioinoculants are a promising solution to improve crop health and resilience, but ensuring their persistence and activation in the field remains a challenge. This study examined how dormant root-zone associated bacteria reactivate after exposure to the plant stress hormones abscisic acid and salicylic acid. The experiment revealed that certain bacteria taxa could reactivate in response to these plant stress signals and persist for at least two weeks. This work advances understanding of the potential cues for reactivation of beneficial plant-associated microbes and supports goals toward developing microbial solutions for sustainable agriculture.

## Introduction

Plant hormones, called *phytohormones*, are essential for a plant’s response to stress. Phytohormones improve germination, upregulate antioxidant systems to reduce reactive oxygen species (ROS) accumulation, induce systemic acquired resistance, and assist in cell signaling.

They are also vital growth regulators that can impact plant metabolism and are often externally applied to plants to improve plant growth under stress conditions (1). Several microbes are known to produce phytohormones or phytohormone mimics (2–5) which influence phytohormone networks in plants and subsequently affect microbial metabolism. Thus, phytohormone regulation in plants is considered to be connected to microbial phytohormone production, together forming inter-organismal systems that can support plants (6).

The plant rhizosphere and proximal root zone are nutrient-rich compartments due to root exudation and thus are heavily colonized by microbes (7, 8). Sugars and amino acids within root exudates serve as nutrients for microbes (9, 10), which in turn can assist plants by improving their nutrient acquisition (11) and protecting against stress (12, 13). Specifically, microbial production of phytohormones (6) such as abscisic acid (ABA), auxins (indole 3-acetic acid), salicylic acid (SA), and cytokinins can improve a plant’s ability to cope with stress. These well-known plant-microbe interactions have been extensively investigated (6, 14–16).

The phytohormones ABA and SA play an important role in plant abiotic stress tolerance. Abscisic acid belongs to a group of phytohormones called sesquiterpenoids, which regulate plant growth. ABA-induced signaling can regulate the expression of stress-responsive genes and contribute to stress tolerance (17). During drought, ABA induces stomatal closure and thus controls transpiration (18). Exogenous ABA application to plant seeds before sowing can protect a drought-sensitive wheat cultivar from drought-induced oxidative damage by increasing the activity of the antioxidant enzyme peroxidase (19). Salicylic acid also modulates antioxidant enzyme activities (20, 21). Similar to ABA, the application of SA to plant tissues has been shown to relieve water stress (22), potentially enabling continued plant growth despite drought (21).

While much is understood about a plant’s coping mechanisms during stress via phytohormone regulation of cellular pathways, we are still learning how phytohormones impact rhizosphere microbiome assembly during stress. It has been shown that SA signaling can increase microbial community diversity and result in the colonization of specific taxa (23–25). Similarly, low molecular weight organic acids such as malic acid, citric acid, and fumaric acid secreted by roots have also been shown to serve as a source of carbon substrate and signaling molecule for rhizobacteria recruitment (26–28). Direct evidence of phytohormones serving as a carbon source for microbes is more limited; however, some studies demonstrated increased soil respiration when treated with one mM abscisic acid (29) and higher soil respiration when soils were treated with root exudates from drought exposed plants compared to exudates from non-droughted plants (30). However, there is indirect evidence that rhizobacteria recruitment can be enhanced in response to specific phytohormone precursors such as L-tryptophan, a precursor for auxin synthesis (31). This study showed that auxin-producing *Pseudomonas fluorescens* stimulated the root growth of radish, which exudes high levels of tryptophan. If microbes use phytohormones as a carbon source, it is possible that they could cause a soil-priming effect that stimulates microbial respiration. This has been shown in experiments where low levels of jasmonic acid (JA) and 1-aminocyclopropane-1-carboxylic acid (ACC) addition to soils caused a relatively large increase in soil respiration (29). Such priming effects have been documented for various organic substrates in root exudates (32–35).

The rhizosphere microbiome members recruited during stress are expected to support a plant’s ability to recover from that stress or exploit the plant during stress. Thus, understanding the role of stress-associated phytohormones in determining rhizosphere microbiome assembly is important for fully understanding the complex dialogue and feedback between a plant and its microbiome. Phytohormone release or leakage into the root zone is also one potential mechanism by which plants may “cry for help” via root exudation to attract supportive microbial partners from the root zone to the rhizosphere (36).

A complicating factor in understanding plant-soil-microbial interactions during stress is the unclear contribution of those soil microbes that reactivate from dormancy. A large fraction of the soil bacterial community can be dormant at a given time (80-90% of cells and 55% of taxa) with dormant cells remaining inactive for prolonged periods (37). Even nutrient-rich rhizosphere soil can have substantial microbial dormancy, with around 40-80% inactive cells reported in the rhizosphere soil of bean plants (38). Thus, we reasoned that the rhizosphere and proximal root-zone compartment may be highly dynamic in activity switching during stress, with some taxa reactivating in response to plant signals (and possibly supporting the plant’s stress response) while others initiate dormancy to avoid the stress by protecting themselves (39).

The overarching objective of this study was to understand how the root-zone bacterial microbiome activates and inactivates in response to phytohormones that signal plant stress. Our null hypothesis was that the microbiome does not react to phytohormone exposure. We had three alternate and non-mutually exclusive hypotheses:

> **H1: The microbiomes of root-zone soils from different plants have different activation and assembly dynamics in response to phytohormones.** The motivation for this hypothesis is our previous work on these plant genotypes (40) and the literature, which shows that different plant species harbor different bacterial microbiome structures. If supported, this hypothesis suggests the importance of the plant’s ability to locally assemble and activate “custom” microbiome members to respond to its own phytohormones.
>
> **H2: Certain bacterial lineages generally activate in response to phytohormones regardless of the plant**. The rationale for this hypothesis is that the phytohormones could serve as resources for microbes. Therefore, the bacteria that can utilize exogenous phytohormones may generally activate if provided either SA or ABA from any plant species. This activation pattern may indicate that taxa respond to resource availability, not a stress-specific “cry for help.”
>
> **H3: There is a hormone-specific activation of certain bacterial taxa**. Each phytohormone indicates a different type of stress for the plant: mainly pathogen infection in the case of SA and drought in the case of ABA. Thus, if some bacterial lineages activate after exposure to one of these phytohormones but not both, it would be consistent with a bacterial taxon’s response to a plant’s “cry for help” given a specific stress.

We addressed these hypotheses using a mesocosm experiment in which we exposed root-zone soils to either a stress-associated phytohormone or controls. We collected root-influenced soil adjacent to plant roots (<20cm) from two different cropping systems: common bean (*Phasoelus vulagris*) and switchgrass (*Panicum virgatum*). These two plants were chosen for their different life strategies (annual and perennial) and families (Fabacaea and Poaceae) and, therefore, to provide a contrast for understanding the potential generalizability of microbiome responses to phytohormones. We used field soils to retain the legacy of the crop in assembling the bacterial communities relevant to agricultural conditions. We applied the stress-associated phytohormones abscisic acid (drought) and salicylic acid (pathogen infection) to the mesocosm soils. We used 16S rRNA and rDNA amplicon sequencing to designate a taxon-level activity state (likely active, likely inactive) before and after phytohormone application. We also related microbiome data to soil nutrient concentrations to understand its impact on the microbial community shifts in activity across treatments and over time.

## Results

### Experimental design

There were four experimental conditions including controls: methanol carrier control (water and methanol), abscisic acid (water, methanol, and abscisic acid), salicylic acid (water, methanol, and salicylic acid), and a water control (water only) and three time points (Day 1, Day 7 and Day 14). Each condition was replicated four times for 16 mesocosms (four conditions, four replicates) for each soil type (See **Figure 1** for experimental design). This design was executed for two soils (bean and switchgrass root zone), resulting in 32 mesocosms. Including the field soil, pre- and post-dry samples, water acclimation, and the time series for each experimental condition, there were 333 microbiome samples across the bean and switchgrass soils.

**Figure 1.**
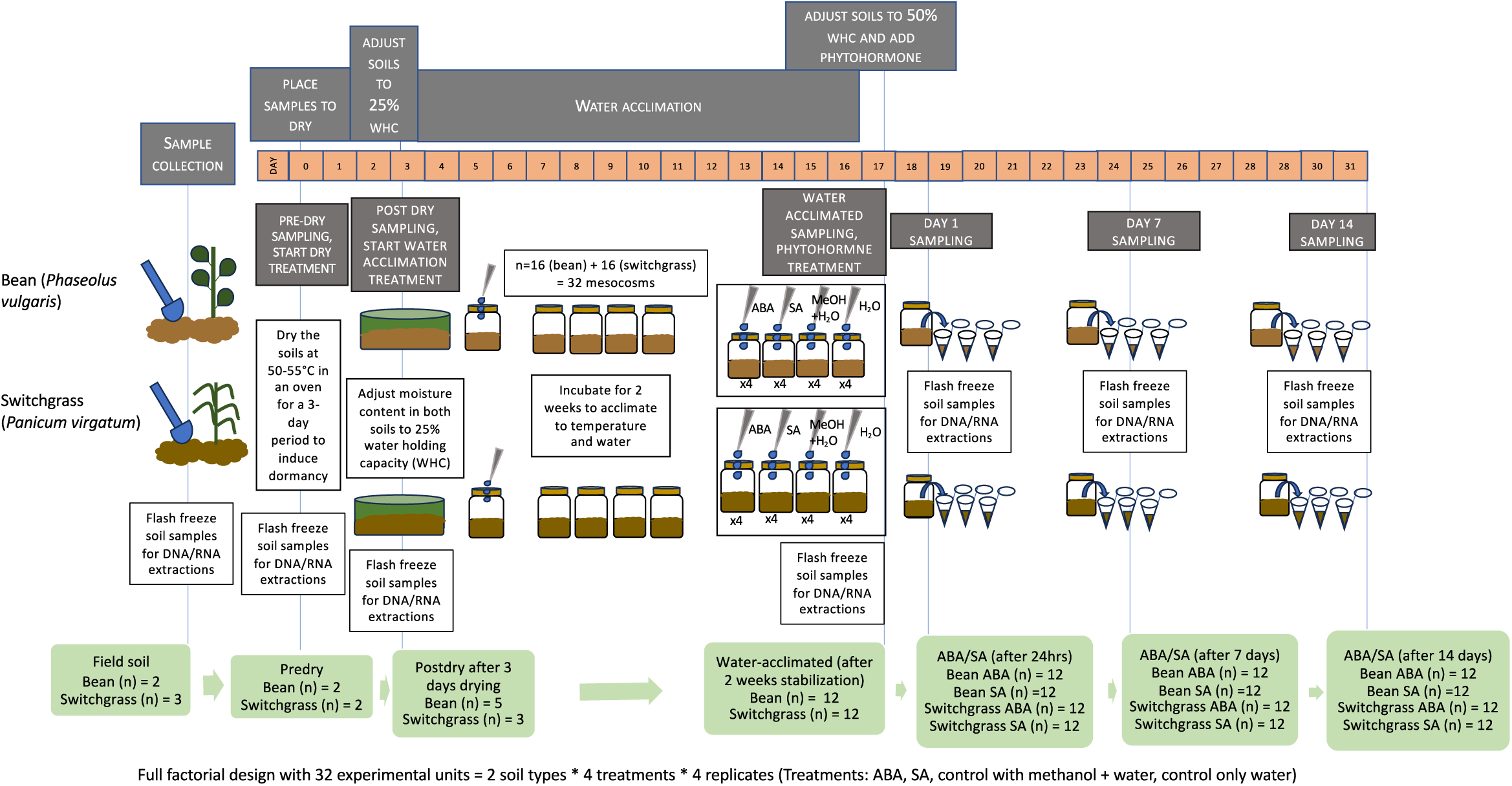
Schematic of the soil mesocosm experimental design and sampling approach to assess how root zone bacterial communities activate and assemble in response to phytohormones that indicate plant stress. The design was a time series of field-collected root zone soils that were first deactivated by drying and then acclimated to water before the experimental addition of phytohormones. The water acclimation step was used to partition the response to moisture increase from the response to the phytohormones. Soils were amended with either a stress-associated phytohormone (salicylic acid, SA, or abscisic acid, ABA) or a methanol carrier or water control. WHC is water holding capacity.

### Sequencing summary

We sequenced a total of 333 DNA samples and 333 cDNA samples from this experiment. After rarefying all read libraries to 12,000 reads per sample (**Figure S1A**), 324 DNA and 324 cDNA samples met the minimum observation effort. After removing all contaminants from the OTU table using package decontam (**Figure S1 B, C**), 13,138 OTUs were used to determine the active community.

### Immediate activation and persistence of the root zone bacterial microbiome to phytohormones

After determining the likely active taxa at Day 1 post-hormone addition, we assessed their relative abundances (DNA counts) before and after phytohormone application. To focus on enrichments observed only in the phytohormone treatments, we also compared the enrichment of specific taxa after hormone addition (Day 1) to the water-acclimated sample and the water and methanol controls. There were statistical differences in the active bacterial community structure between the water-acclimated and Day 1 for bean ABA soil (PERMANOVA F=3.95, R2=0.15, p=0.002), bean SA soil (PERMANOVA F=3.96, R2=0.15, p=0.006), switchgrass ABA soil (PERMANOVA F=10.50, R2=0.32, p=0.001) and switchgrass SA soil (PERMANOVA F=2.31, R2=0.09, p=0.002).

In the ABA-treated soils, three OTUs were enriched by log two-fold at Day 1 compared to the water-acclimated sample and not enriched in water and methanol controls (**Figure S2A**). These included Actinobacteriota OTUs from Family Myxococcaceae, genus *Microbispora*, and genus *Streptomyces*. However, these enrichments were detected only in the switchgrass soils, and there were no OTUs enriched in bean soils treated with ABA on Day 1. Furthermore, all three taxa enriched in the switchgrass ABA soils switched from either an inactive or below detection state in the water-acclimated sample to high abundance at Day 1 after hormone addition (**Figure S3B**).

Similarly, for the SA treatments, the two OTUs belonging to genus *Microbispora* were log two-fold enriched at Day 1 compared to the water acclimated sample in both bean and switchgrass and were enriched solely in the SA treatments to the exclusion of the methanol and water controls (**Figure S2B**). Furthermore, the response pattern suggested a likely switch from inactive (detected in DNA but not RNA) to active for both taxa. One was enriched on Day 1 in bean soils, and the other OTU was enriched on Day 1 in both bean and switchgrass (**Figure S3 C, D**)

Several taxa belonging to classes Myxococcia, Actinobacteria, Alphaproteobacteria, and Bacilli were resuscitated within one day after SA addition to the bean soil (**Figure S3C**), but most of these taxa were also responsive to methanol (**Figure S3E**). Two Actinobacterial OTUs (both belonging to the genus *Microbispora*) were activated in response to SA treatment in bean soils and not in the methanol controls. These two OTUs were log two-fold enriched after Day 1 of SA exposure compared to the water-acclimated soils. However, only one of these *Microbispora* OTUs was represented in the top 50 abundant and active taxa for the bean SA treatment but both are included for comparison (**Figure S3C**). Both taxa were detected as active in methanol controls but not as consistently enriched as in the SA treatment. However, apart from the log two-fold enrichments mentioned above, no substantial response to ABA or SA was noted among other active OTUs for switchgrass beyond their similar response to the methanol on Day 1.

While several active taxa increased abundances after phytohormone addition (**Figure 2**), many of these were also active in the methanol controls (**Figure S4**). However, some active taxa persisted past Day 1, a pattern consistent with growth if the phytohormones were used as a resource. We observed the persistence of active taxa over 14 days for four of the enriched taxa (indicated in red in **Figure 2**), including a *Microbispora* for switchgrass and bean SA and switchgrass ABA and a Myxococcaceae and *Streptomyces* in switchgrass ABA. Notably, persistent activation over time could also be an indirect response to the phytohormone addition (e.g., due to priming), which would not be distinguishable here.

**Figure 2.**
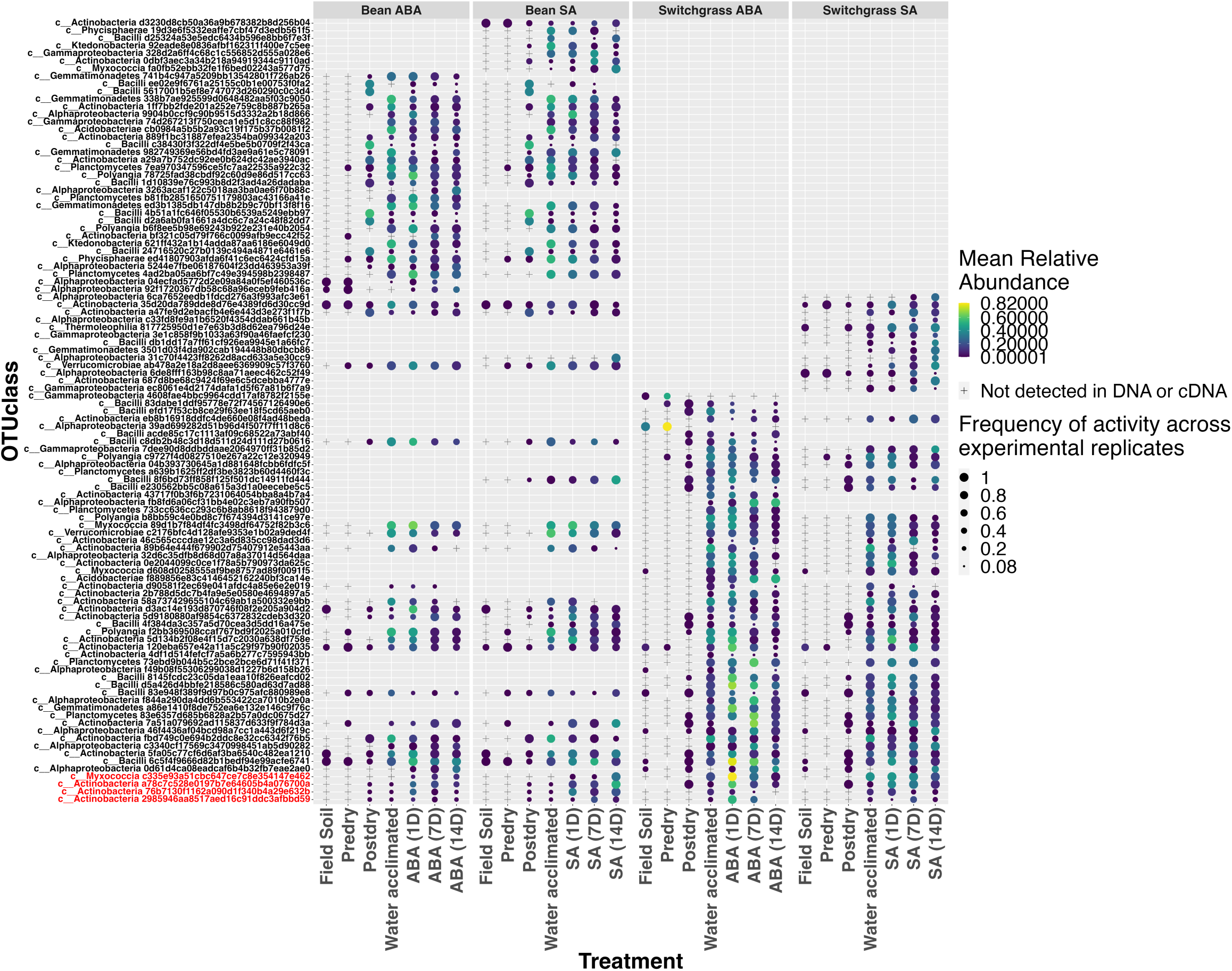
Bacterial microbiome community activation, persistence, and assembly before and after phytohormone addition to bean and switchgrass root zone soil. The relative abundances of active taxa that were log two-fold enriched (DESeq analysis) at Day 1 as compared to the water-acclimated sample are shown in red. All other taxa include the 50 most abundant active members in each soil/phytohormone condition. All responses shown were specific to a phytohormone exposure (salicylic acid, SA, or abscisic acid, ABA) and were not observed in controls. Inactive taxa have a relative abundance of zero, so they do not contain a symbol on the graph and are left blank for that time point/treatment. Taxa not detected in either DNA or cDNA data set are denoted by a + shape. Color gradient denotes increased relative abundances as color changes from blue to yellow. The shape size of each bubble denotes the consistency of detection across experimental and technical replicates of soil mesocosms (e.g. with 1 denoting activity in all replicates and 0.5 denoting activity in half). Empty rows indicate that the taxon was not detected within the top 50 most abundant members for that condition. Empty spaces in rows that also have symbols indicate that the taxon was not classified as active within those samples (columns) but was detected in DNA.

### Addressing H1: The microbiomes of root-zone soils from different plants have different activation and assembly dynamics in response to phytohormones

Bean and switchgrass soils differed distinctly in their active community structure across all time points and treatments (PERMANOVA F=29.05, R2=0.08, p=0.001). For the ABA treatment, there was a higher proportion of Bacilli and Verrucomicrobiae in bean soils on Day 1 after ABA addition, but a higher proportion of Actinobacteria and Myxococcia in switchgrass soils. Then, Alphaproteobacteria increased proportion 7 and 14 days after ABA exposure in switchgrass compared to bean soils (**Figure 3**). For the SA treatment, there was also a relatively higher abundance of active Actinobacteria on Day 1 in switchgrass soils than in bean soils. However, Actinobacteria declined in switchgrass soils over time and was generally replaced with Alphaproteobacteria. In bean soils, active Actinobacteria increased over time, with a concurrent relative decline in several other bacterial classes (**Figure 3**).

**Figure 3.**
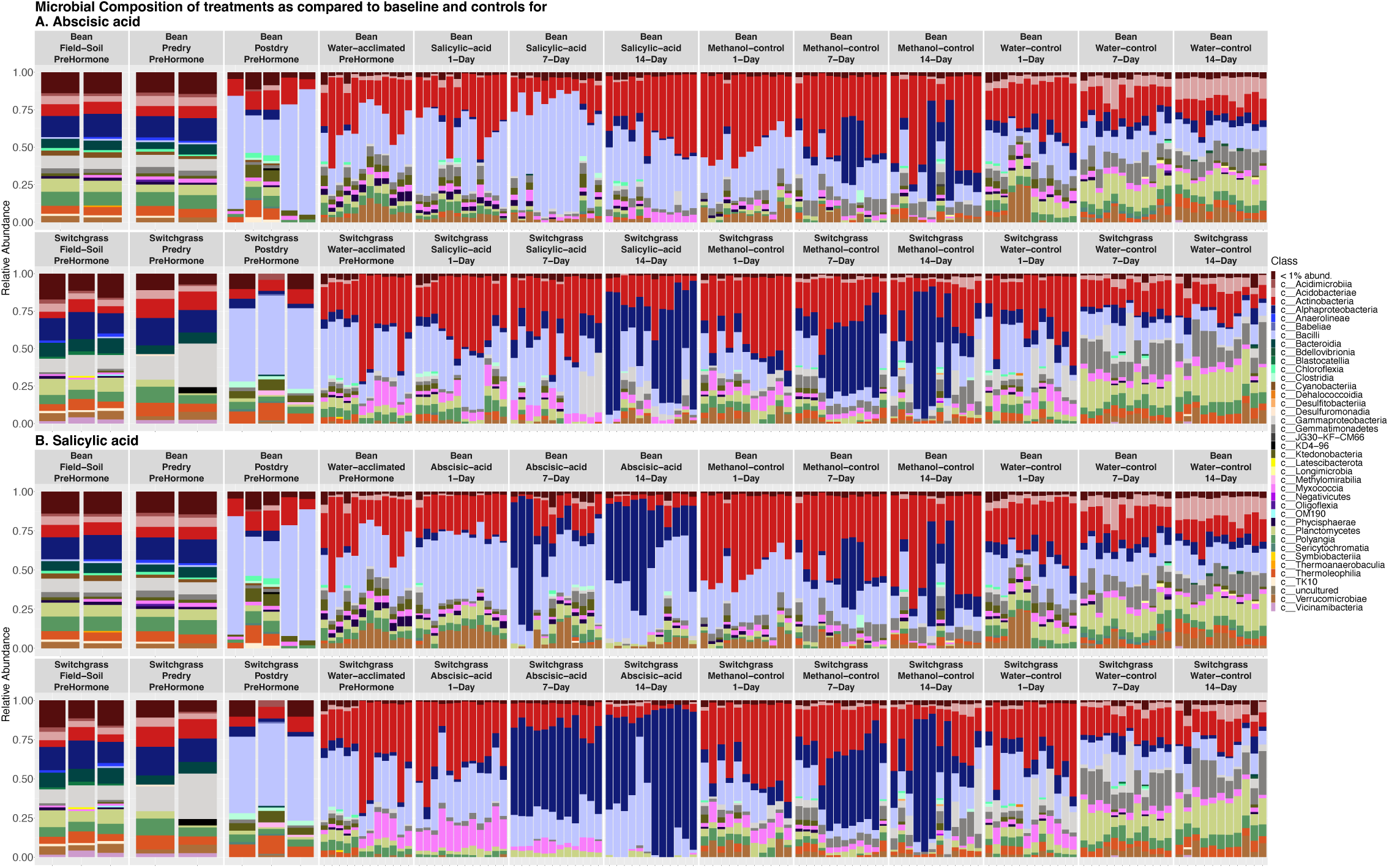
Active bacterial community structures differed between bean and switchgrass. Barplots show broad changes in the bacterial community composition over time and across treatments in bean and switchgrass root zone soils for **A)** Abscisic acid and **B)** Salicylic acid.

Several taxa were enriched on Day 1 after ABA addition in switchgrass soils compared to the water-acclimated control. Specifically, OTUs belonging to phyla Actinobacteriota and Myxococcota were enriched by log two-fold on Day 1 compared to the pre-hormone time point (**Figure S2**). Bean soil, however, showed no taxa enriched on Day 1 in response to ABA. For SA treatment, one OTU belonging to the Microbispora was enriched on Day 1 in both bean and switchgrass. A different *Microbispora* OTU was also specifically enriched in bean soil in response to SA treatment (**Figure S2)**.

### Addressing H2: Certain bacterial lineages generally activate in response to phytohormones regardless of the plant

One genus *Microbispora* OTU, was enriched in response to both ABA and SA in switchgrass soils on Day 1 compared to the water-acclimated sample. This same OTU was also enriched in bean soils in response to SA. Thus, this lineage responded consistently across different soils and phytohormones.

We also related changes in the active bacterial community to measured soil properties to consider any co-occurring factors (**Figure S5).** We found that consistent phytohormone responses by microbial taxa were unlikely to be attributable to concurrent pH or organic matter changes because the treatments generally had different properties (**Figure S5**). Nitrate (in bean) and ammonium (in switchgrass) explained the variation in the soil communities after receiving water (**Figure S5**). Overall, soil differences in pH, nitrate concentrations, and organic matter content explained about 27% of the variation in the active bacterial community in bean soils. In contrast, pH, ammonium concentration, and organic matter content explained about 22% of the variation in the active bacterial community in switchgrass.

### Addressing H3: There is a hormone-specific activation of certain bacterial taxa

We found no strong evidence of any taxa common to both plants’ root zone soils and similarly responsive to either ABA or SA exposure, and thus, H3 was not supported. Two OTUs (family Myxococcaceae and genus *Streptomyces*) were enriched solely in response to ABA in switchgrass. Similarly, one *Microbispora* OTU in bean soil was exclusively enriched in response to SA, indicating a hormone-specific and soil-specific response from these taxa.

However, there was a generally consistent response dynamic of the active bacterial microbiome’s structure (beta diversity) to phytohormone addition regardless of the soil origin (**Figure 4, Table 2**). Phytohormone treatment and time both explained global variation in the active community structure. When excluding the three baseline samples (field, pre-dry, post-dry), we found that bacterial community richness was statistically different between crop type (ANOVA F=36.73, SumSq=46434, p<0.001) and time point (water acclimated, Day 1, Day 7 and Day 14) (ANOVA F=8.22, SumSq=31165, p<0.001). In contrast, Inverse Simpson’s diversity, which accounts for richness and evenness, differed only by time point (ANOVA F=3.04, SumSq=35840, p=0.03). There was also a strong interaction between crop and treatment (abscisic acid, salicylic acid, methanol control, and water control) (ANOVA F=12.78, SumSq= 32290, p<0.001). At the same time, treatment strongly influenced Inverse Simpson’s diversity (ANOVA F=4.81, SumSq=47335, p=0.003). Overall, we found that richness decreased over time for abscisic and salicylic acid treatments across both soils (**Figure S6**).

**Figure 4:**
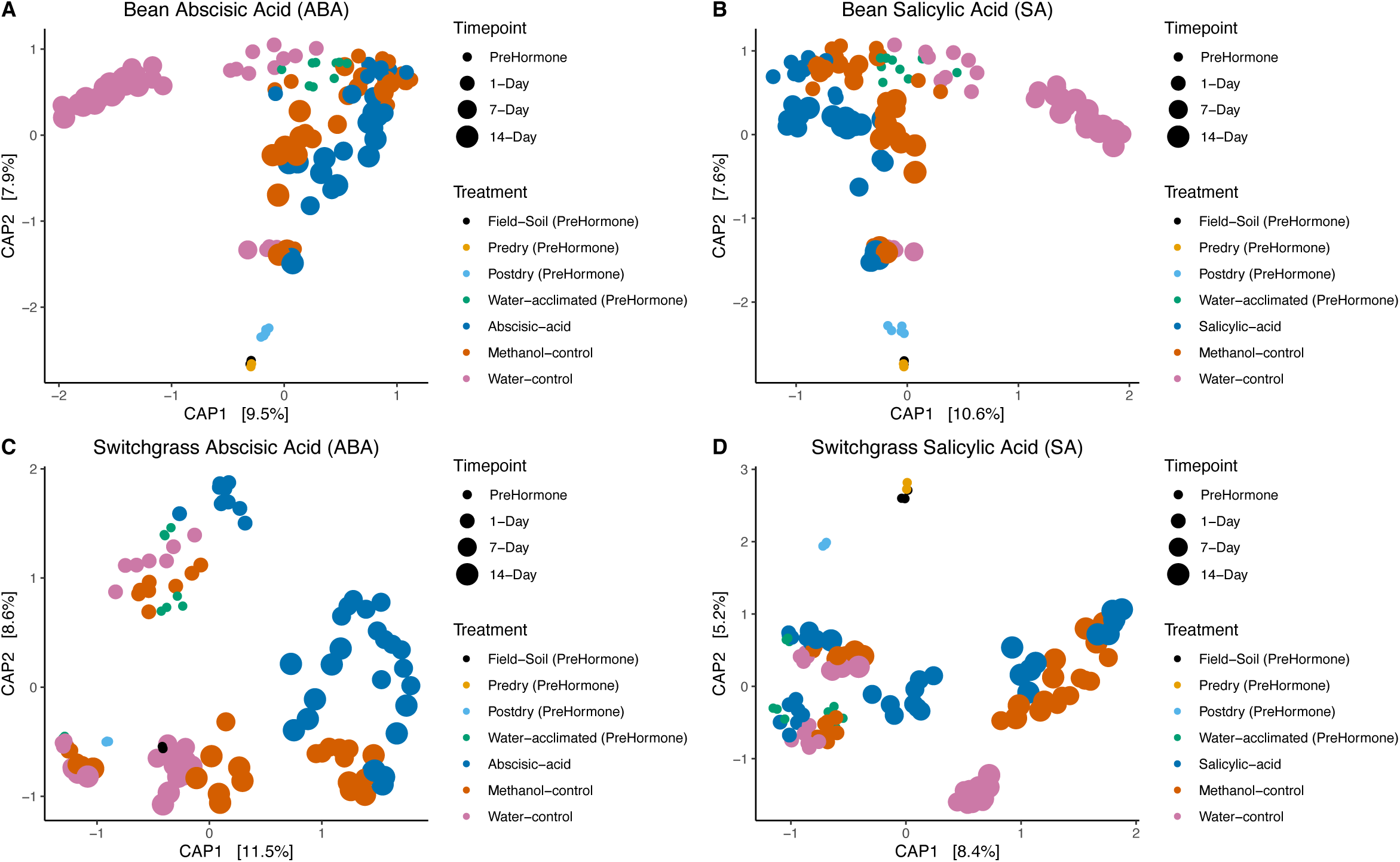
Active microbiome community structure differs across time points for A) Bean soil amended with abscisic acid (ABA), B) Bean soil amended with salicylic acid (SA), C) Switchgrass soil amended with abscisic acid (ABA) and D) Switchgrass soil amended with salicylic acid (SA) treatments. Ordination shows a constrained analysis of principal coordinates (CAP) after excluding the variance attributed to mesocosm. The increasing symbol sizes denote sampling time and the colors denote the treatments.

## Discussion

This study assessed the activation and assembly dynamics of activated root zone bacterial taxa in response to phytohormones produced by plants during stress. Thus, this study provides new insights into which plant-associated bacterial populations immediately activate in response to SA and ABA phytohormones, suggesting that these molecules could serve as reactivation signals. Furthermore, this study also distinguished general and specific activation dynamics of bacterial taxa for two different plants’ root zone soils over two weeks after exposure, allowing improved precision on the ecological underpinnings of these responses. Ultimately, we can weigh the evidence that the activated taxa responded to the phytohormones as a “cry for help” from the plant.

In this study, crop-specific responses of the root zone microbiome to phytohormones indicated a plant legacy on its associated soil that partially determines its microbiome’s response, providing evidence in support of H1. Given that most plant species harbor unique microbial communities (41–43), it is not unreasonable that each plant species may have a different microbiome response to stress indicators like phytohormones. While switchgrass responded to ABA via immediate enrichment of three OTUs within 24 hours of phytohormone application, bean soils show no enrichment of any taxa in response to ABA. Similarly, one Actinobacterial OTU was enriched in response to SA treatment only in bean but not in switchgrass soils, suggesting a legacy effect that impacts the recruitment of active members to the bean rhizobiome. This finding agrees with the results observed in our previous study that assessed bacterial reactivation after drought (40). Other studies have also shown that root exudate-induced soil respiration can be observed nearly immediately within 6-21 h (30).

Actinobacterial genera are widely reported to be associated with drought stress in plants (44), so it was not surprising that we detected several Actinobacterial taxa that activated in response to ABA and SA exposure, showing support for H2 among these lineages. Of the most abundant active members in bean soils exposed to ABA and SA, we found an Alphaproteobacterial OTU (*Microvirga*) that activated after hormone addition within 24 hours but did not activate in the methanol control mesocosms within the same time frame. This Alphaproteobacterial genus *Microvirga* has also been shown to alleviate drought stress in plants such as cowpeas, such that cow peas inoculated with *M. vignae* had no differences between water deficit and well-watered conditions concerning stomatal conductance, shot dry mass, or N accumulation (45). This same study found that ABA biosynthesis and ABA-dependent genes were less upregulated under drought when plants were inoculated with a *Microvirga sp*. compared to a *Bradyrhizobium sp*, suggesting that the presence of *Microvirga* relatively decreased the plant’s typical ABA production during stress.

We found a specific Actinobacterial genus, *Microbispora,* that consistently responded to both ABA and SA across bean and switchgrass. This response pattern indicates that this microbial group could be a generic responder to phytohormones as stress signals. Actinobacteria have been widely explored as targets for developing bioinoculants. *Microbispora* and *Streptomyces* have been suggested to be both soil-inhabiting and endophytic genera, though only a limited number of species are likely to have this dual habitat preference (46). *Microbispora* and other Actinobacterial species belonging to *Actinoplanes, Streptomyces, Rhodococcus*, and *Micromonospora* have also been explored for direct/indirect plant growth-promoting capabilities via secretion of phytohormones, siderophores, and antimicrobial agents (47, 48).

Across the two soils used in this study (bean, switchgrass), we found no taxa responding consistently to either ABA or SA, indicating that hormone-specific responses may not be conserved across different plants, so H3 was not supported. However, we did find hormone-specific responses to either ABA or SA within each soil type. For example, we found one Actinobacterial species (*Microbispora*) that uniquely responded in bean to SA, while one Actinobacterial (*Streptomyces*) and one Myxoccoccal OTU responded uniquely to ABA in switchgrass. While we found signatures of Gram-positive bacteria activating to phytohormones, other studies have found a declining abundance of Gram-positive biomarkers following phytohormone addition (29). However, these other studies did not discriminate activity states but rather assessed the microbial communities using phospholipid-derived fatty acid profiles, and a negative response to phytohormones also may be due to an investment in complementary survival strategies such as osmolyte production and spore production as opposed to growth and turnover (49–51).

The literature shows that phytohormone signaling can play a major role in rhizobiome assembly (25). For instance, using natural selection T. Kalachova et al. (52) showed a reduction in disease severity of Arabidopsis plants infected with foliar pathogen *Pseudomonas syringae*. This was associated with SA-mediated induction of defense mechanisms and a shift in the soil bacterial community, but not the fungal. While one study showed that ABA and polyacrylamide treatments to soil coupled with foliar application of ABA reduced the relative abundances of Actinobacteria in the rhizosphere using amplicon DNA sequencing and increased drought resistance in forage grass (53), other studies have shown that Actinobacteria increase in plant roots and rhizosphere soil during drought likely due to expansion of C allocation in root exudates (44).

Furthermore, certain Actinobacteria, such as *Streptomyces,* have been shown to increase the ABA content of wheat leaves to upregulate the expression of drought-resistance genes (54). Thus, there could be implications for a positive feedback loop in Actinobacterial responses to a plant’s cry for help. The role of ABA as a carbon substrate for Actinobacteria is also indicated. For example, *Rhodococcus* sp. utilized ABA as the sole carbon source in batch culture (55). These findings support the idea that phytohormone substrates can be used to select rhizosphere microbes (56) and corroborate our finding of active Actinobacterial in this study. Overall, there are limited experiments that consider microbiome reactivation with plant signaling.

A limitation of this study is that we did not assess whether microbes utilized the phytohormones ABA and SA. A previous study reported that increasing ABA concentrations caused increased soil respiration, suggesting microbial utilization of ABA as a substrate. However, they also observed a disproportionate increase in respiration relative to the minimal phytohormone inputs of JA and ACC, suggesting that these instead stimulated microbial mineralization of existing soil carbon (29). Here, we observed a continued persistence of activated taxa in response to phytohormones over 14 days of a few taxa (specifically the log two-fold enriched taxa in red – **Figure 2**), which could be an indirect response due to a potential priming effect. However, the immediate activation of bacterial taxa within 24 hours suggests that the phytohormones can dually serve as a signal, potentially pointing to a plant’s cry for help. Another limitation of this study is that the 16S rRNA and rDNA sequencing method provides a population-level average of whether a taxon is active (yes or no) and no indication of relative activity levels. Thus, we cannot know whether all individuals within that taxon consistently responded or if some cells remained inactive while others became disproportionately active.

Overall, this experiment’s results show that microbial reactivation can be used to identify immediate and then persistent bacterial responders among the root zone soil microbiome to plant stress signals. The Actinobacterial members, such as *Microbispora* sp., were consistently noted among the responsive bacterial populations across the different phytohormones and soils. The next steps should investigate the molecular mechanisms and the dynamic host-microbiome feedback with *Microbispora* sp. given phytohormones as stress signals from the plant.

## Methods

### Soil collection, processing, and storage

Five kilograms of root-zone soil was collected from four replicated switchgrass plots at Kellogg Biological Station’s Great Lakes Bioenergy Research Center Bioenergy Cropping Systems Experiment in November 2019. Approximately 4 kg of soil was also collected from four soil core replicates for common bean (*Phaseolus vulgaris* L.) from a rotation plot at the Michigan State University’s Agronomy Farm in East Lansing, also in November 2019. Approximately 1000 g of soil from each plot was collected from the top 10 centimeters for each replicate using an ethanol-sterilized soil auger of 10 cm diameter and placed in Whirlpak bags. Soils were transported to the lab in a cooler with ice and then stored at 4°C until further processing.

The following day after sample collection, soils from replicate plots were composited into ethanol-sterilized buckets and hand-mixed to homogenize. Composited soils were sieved through 4mm mesh to remove roots and debris. Three 0.5g subsamples of each composited soil were flash-frozen in liquid nitrogen for active microbiome assessment and designated as the “field-soil” samples. The remaining sieved soil was stored at 4°C until the start of the experiment. All flash-frozen soil samples were stored at −80°C.

Soil organic matter, pH, lime index, phosphorous, potassium, calcium, magnesium, nitrate, and ammonium levels of the “field-soil” soils were analyzed by the Michigan State Soil Plant and Nutrient Laboratory according to their standard protocols listed here https://www.canr.msu.edu/fertrec/about/detailed-instructions. (**Table 1**).

**Table 1.** Nutrient analysis of bean and switchgrass root zone soil before the experiment (pre-dry sample). P-Phosphorus, K-Potassium, Ca-Calcium, Mg-Magnesium, OM-organic matter, NO_3_-N-Nitrate-N, NH_4_-N-Nitrate.

| Soil | pH | Lime | BrayP1 P | K | Ca | Mg | OM | NO<sub>3</sub>-N | NH<sub>4</sub>-N |
| --- | --- | --- | --- | --- | --- | --- | --- | --- | --- |
|  |  | Index | ppm | ppm | ppm | ppm | % | ppm | ppm |
| Switchgrass | 6.0 | 69 | 27 | 96 | 930 | 146 | 2.5 | 1.5 | 2.6 |
| Common bean | 6.2 | 69 | 97 | 151 | 486 | 82 | 1.9 | 0.7 | 1.6 |

**Table 2.** Permuted analysis of variance (PERMANOVA) to assess global differences in active microbial community structure across experimental treatments and time points.

|  | Degrees<br>of<br>Freedom | R2 | Pseudo-F value | P value | Degrees<br>of<br>Freedom | R2 | Pseudo-F<br>value | P value |
| --- | --- | --- | --- | --- | --- | --- | --- | --- |
|  |  | Bean |  |  |  | Switchgrass |  |  |
| Treatment | 4 | 0.19 | 10.28 | 0.001 | 4 | 0.16 | 9.24 | 0.001 |
| Time point | 2 | 0.06 | 6.63 | 0.001 | 2 | 0.11 | 12.79 | 0.001 |
| Treatment: Time<br>point | 6 | 0.10 | 3.66 | 0.001 | 6 | 0.11 | 3.98 | 0.001 |

Gravimetric soil moisture was determined by comparing the mass difference between the initial field soil sample and a sample dried at 92°C for 4 days. Water holding capacity (WHC) was assessed by taking 10g of dry soil and placing it in a funnel lined with filter paper (57). The dry soil was saturated with water and allowed to drain for 2 hours. The mass of the original soil, the mass of water added, and the mass of water drained were used to calculate WHC as percent gravimetric soil moisture. Before setting up the mesocosms, 0.5 g of stored soil was flash-frozen denoting “pre-dry” condition accounting for any shifts in microbial community due to storage, thus capturing a true baseline reference community. The first step of the experiment was drying the soils to induce dormancy and use it as a reference for subsequent treatments. Soils were dried in a 55°C incubator for 3 days and after drying, soil subsamples were flash-frozen as the “post-dry” treatment before adding water and phytohormones.

### Mesocosm set-up, phytohormone treatment, and maintenance

To create each mesocosm, 100g dried soil was adjusted to 25% WHC and placed into 473 mL sterile glass jars (**Figure 1**). Mesocosms were maintained at 27°C with lids secured loosely to ensure aerobic conditions. Mesocosm manipulations and samplings were performed inside a sterilized biosafety cabinet. Every 2-3 days, mesocosms were massed, and any water loss was replaced with sterile water to maintain soil moisture.

Before hormone addition, mesocosms were incubated for two weeks to acclimatize to the baseline water content of 25% WHC. This water acclimatization step was included to ensure that changes in activation could be attributed to the experimental treatment of hormone addition and not exclusively to water addition from a dry state, as it has been well-reported that soil microorganisms activate in response to water (58–61) After two weeks, three 0.5-g subsamples from four replicate mesocosms designated as “post 25% WHC” were flash frozen in liquid nitrogen and designated as “water acclimated” samples. After sample collection at two weeks, the “post 25%” mesocosms were supplemented with the required water to reach 50% WHC. They were designated as “water control” samples, while the rest of the mesocosms received phytohormones dissolved in methanol and methanol alone (carrier control) mixed with enough water to reach 50% WHC.

Phytohormones were dissolved in methanol to create a 0.3 M solution, and 1 mL of the phytohormone-methanol solution was filter sterilized (0.2 micron) before adding to the mesocosms in combination with enough water to achieve 50% WHC. We normalized the hormone addition based on the amount of carbon added per g of dry soil, and a 3M solution for ABA and SA resulted in approximately 250-500 ug C g-1 dry soil for SA and ABA. Phytohormone concentrations reported in soil are typically in the nanomolar range (62). However, we chose a concentration that was 4-7 times the reported values used in the literature (which ranges anywhere between 5-64 ug C g-1 dry soil (30, 63)), to elicit a measurable response to the phytohormone such that comparisons across treatments would be possible. Methanol carrier controls received 1 ml of methanol supplemented with water to reach 50% WHC. After adding the aqueous solution, mesocosm soil was mixed using a sterile spatula. The soil was non-destructively sampled from each mesocosm and immediately flash-frozen on days 1, 7, and 14 after hormone addition.

Thus, there were four conditions in total – water control, methanol control, ABA, and SA. With four replicate mesocosms for four treatments and three time points collected in three replicates, n = 144 total samples for each soil type after water acclimation, totaling 288 samples across both soils.

### RNA/DNA coextractions

A manual phenol-chloroform nucleic acid extraction was performed to obtain RNA and DNA from the same cell lysis pool (64) with minor modifications. For the modifications, we used Qiagen Powerbead Garnet Tube (0.70mm), and after a 30 s lysis in a bead beater, we used a 10-minute centrifugation at 12,000 g at 4°C. After adding the chloroform-isoamyl alcohol (24:1), the tubes were inverted several times to form an emulsion, followed by a 5-minute centrifugation at 16,000 g at 4°C. After adding 30% PEG6000-1.6 M NaCl, the tubes were inverted several times to mix the contents and then placed on ice for a 2-hour incubation. After incubation, the samples were centrifuged for 20 minutes at 16,000 g at 4°C. After centrifugation, the supernatant was removed, and 1.0mL of 70% EtOH (stored at −20°C overnight) was added. The samples were centrifuged for 20 minutes at 16,000g at 4°C, and the ethanol wash was removed. The samples were centrifuged for an additional 20 seconds. Residual ethanol was removed with a pipette, and the pellet was left to air-dry before they were resuspended in 30 _μ_L of nuclease-free water. Negative extraction controls (tubes with reagents and beads) were processed alongside experimental samples. Nucleic acids were visualized using agarose gel electrophoresis and validated with a band for DNA and RNA (64). All DNA and RNA samples were quantified using a Qubit® dsDNA BR assay kit and RNA HS assay kit on a Qubit 2.0 fluorometer (Invitrogen, Carlsbad, CA, USA). Samples were stored at −80°C.

### DNase treatment and cDNA synthesis

RNA samples were purified using an Invitrogen TURBO DNA-free™ Kit with a slightly modified protocol. A six _μ_L aliquot of the nucleic acid coextraction was treated with one _μ_L of 10X TURBO DNase Buffer and three _μ_L of Turbo DNase enzyme and then incubated at 37°C for 30 minutes. After this incubation period, two _μ_L of DNA inactivation reagent was added to each tube before incubating at room temperature for five minutes. Then, the sample was centrifuged at 2,000 g for five minutes at room temperature. The purified RNA was used as a template to form cDNA using Invitrogen’s SuperScript™ III First-Strand Synthesis System using random hexamers per kit directions and with negative controls to assess for reagent contamination. Final cDNA samples were stored at −80°C.

### PCR check and gel electrophoresis

16S rRNA gene V4 PCR was performed for every DNA and cDNA sample to ensure the presence of the 16S rRNA gene and the RNA sample to ensure effective DNase treatment. The PCR reaction used a 2X GoTaq Green Master Mix and the primers 515F (GTGYCAGCMGCCGCGGTAA) and 806R (GGACTACNVGGGTWTCTAAT). PCR cycle parameters were: 95°C for 3 minutes; 30 cycles of 95°C for 45 seconds, 50°C for 60 seconds, 72°C for 90 seconds; and a final step at 72°C for 10 minutes. The PCR product was examined via gel electrophoresis. All PCR reactions included a no-template negative control and an *E. coli* DNA template as the positive control.

### Illumina sequencing of the 16SrRNA gene and 16SrRNA

DNA and cDNA samples were sent for 16S rRNA gene V4 amplicon sequencing at the Genomics Research Technology Support Facility at Michigan State University. The V4 hypervariable region of the 16S rRNA gene was amplified using dual-indexed Illumina compatible primers 515f/806r, as described by J. J. Kozich et al. (65). PCR products were batch normalized using an Invitrogen SequalPrep DNA Normalization Plate, and the normalized products recovered were pooled. The pool was cleaned and concentrated using a QIAquick PCR Purification column followed by AMPureXP magnetic beads; it was then quality controlled and quantified using a combination of Qubit dsDNA HS, Agilent 4200 TapeStation HS DNA1000, and Invitrogen Collibri Library Quantification qPCR assays. The pool was loaded onto an Illumina MiSeq v2 standard flow cell, and sequencing was performed in a 2×250bp paired-end format using a MiSeq v2 500 cycle reagent cartridge. Base calling was done by Illumina Real Time Analysis (RTA) v1.18.54, and the output of RTA was demultiplexed and converted to FastQ format with Illumina Bcl2fastq v2.20.0.

Sequence data were analyzed using QIIME2 (66). Briefly, all paired-end sequences with quality scores were compressed and denoised using the DADA2 plugin (67). The denoising step dereplicated sequences, filtered chimeras, and merged paired-end reads. The truncation parameters for DADA2 were determined using FIGARO (68) developed by Zymo Research Corporation. All truncation was performed from the 3’ end for consistent final read lengths. The DNA and cDNA datasets were separately quality controlled and denoised because the additional cDNA amplification step could have introduced different errors for the cDNA than for the DNA amplicons. The quality-controlled DNA and cDNA count tables were merged into a single QIIME2 artifact using the feature-table merge command. Similarly, the DNA and cDNA representative sequences were merged into a single QIIME2 artifact using the feature-table merge-seqs command. The representative sequences from the combined count tables were clustered at 99% identity de-novo, and the clustered representative sequences were classified using SILVA v138 (69) to generate the taxonomy file. Again, 99% sequence identity was used to conservatively account for possible errors in the cDNA amplicons resulting from the additional amplification (RT-PCR). The resulting OTU table and taxonomy files were exported to R for ecological analysis.

### Designating the active bacterial populations

All downstream analyses were performed in R version 4.0. The R package decontam (70) was used to determine the number and identity of contaminants in the dataset and remove them using the prevalence method. Contaminating taxa, mitochondria, and chloroplast sequences were filtered from the datasets. A subsampling depth of 12,000 reads per sample was selected based on rarefaction curves. After subsampling, 16S rRNA to rRNA gene ratios were computed from the cDNA and DNA datasets described in A. W. Bowsher et al. (38). We chose the method that applied a 16S rRNA:rRNA gene ratio threshold >=1. We also included “phantom taxa” and changed all DNA counts = 0 corresponding to RNA counts >0 to DNA counts =1. The DNA counts table was then filtered to include only taxa meeting the criteria to be likely active at the population average (e.g., 16S rRNA:rRNA >= 1). The filtered DNA OTU table was used for all ecological analyses, and we refer to this as the “active community.”

### Ecological statistics

Microbiome analysis was performed in R using packages phyloseq (71) and vegan (72). Permutational analysis of variance (PERMANOVA) was used for multivariate statistics. For data visualization, we used a constrained analysis of principal coordinates (CAP) and partitioned out variances contributed by selected variables. Specifically, switchgrass had a significant mesocosm effect (PERMANOVA F=2.84, R2=0.05, p=0.002), so to account for that, we partitioned out the mesocosm variance using a constrained ordination. We also used soil chemistry data to conduct a constrained analysis of principal coordinates to understand how soil nutrient status affected microbiome community changes across treatments using package phyloseq. Edaphic factors included in the final models were selected with function ordistep from the vegan package with the initial full model incorporating pH, percent soil organic matter, nitrate, and ammonium concentrations. We used differential expression analysis (DESeq2, (73)) to assess the log two-fold changes between treatments and time points. We looked at each taxa’s relative abundance separately to determine the reactivation of taxa from an inactive state. Activity dynamics were evaluated using the methods described in S. Bandopadhyay et al. (40). In this study, we did not set a threshold for detecting phantom taxa across samples. To generate the heatmaps and bubble plots looking at relative abundances for each taxon, we used a max standardization approach using the decostand function in the vegan package in R (72). For alpha diversity, we used the estimate_richness function in package phyloseq with ANOVA type III results reported using R package car (74).

## Data and code availability statement

Sequence data is submitted to the Sequence Read Archive under BioProject PRJNA932434. The scripts to analyze the data and generate figures are on GitHub (https://github.com/ShadeLab/PAPER_Dormancy_resuscitation_phytohormone_mesocosm_Bandopadhyay2025.git.)

## Acknowledgments

Support for this research was provided by the United States National Science Foundation under Grant No. MCB #1817377 to A.S. Additional support was provided by the Great Lakes Bioenergy Research Center, U.S. Department of Energy, Office of Science, Office of Biological and Environmental Research under Award Number DE-SC0018409 and by the National Science Foundation Long-Term Ecological Research Program (DEB #1832042). Additional support was provided by the Michigan State University Plant Resilience Institute by, the USDA National Institute of Food and Agriculture and Michigan State University AgBioResearch. AS acknowledges project support from the European Union (ERC, MicroRescue, 101087042). Views and opinions expressed are however those of the author(s) only and do not necessarily reflect those of the European Union or the European Research Council. Neither the European Union nor the granting authority can be held responsible for them.

## Notes

https://github.com/ShadeLab/PAPER_Dormancy_resuscitation_phytohormone_mesocosm_Bandopadhyay2025/tree/main

https://www.ncbi.nlm.nih.gov/bioproject/PRJNA932434/

### Competing Interest Statement

The authors have declared no competing interest.

### Summary of Updates

Added funding information Added supplemental information file Added URLs to GitHub (analysis) and NCBI (raw sequence data files)

