## Supplemental Information for "Activation dynamics and assembly of root zone soil bacterial communities in response to stress-associated phytohormones"

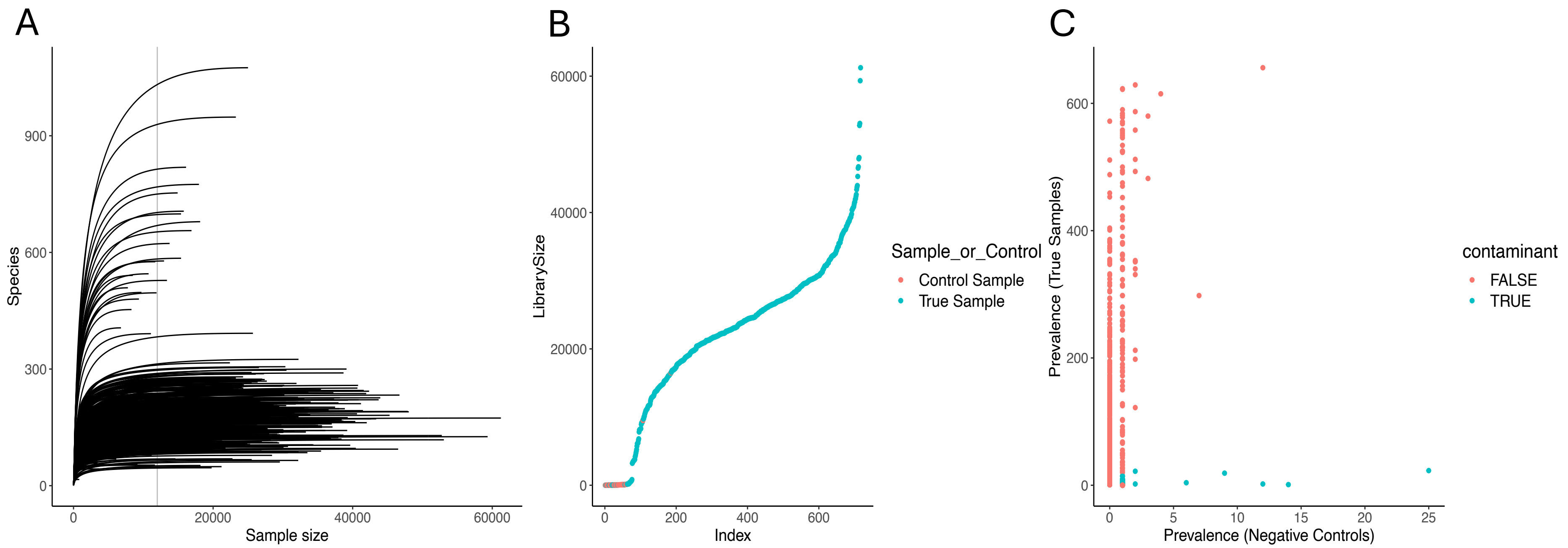


**Figure S1**. **Sequencing quality assessments. A)** Rarefaction curves of 16S rRNA gene amplicons for DNA and cDNA, based on species units, which are operational taxonomic units, OTUs, defined at 99% sequence identity of amplicon reads. Vertical gray line indicates sampling depth (12,000 reads) to which sequences were rarefied for analyses. **B)** Library size of samples. **C**) Identification of contaminants via a prevalence method using package decontam in R.

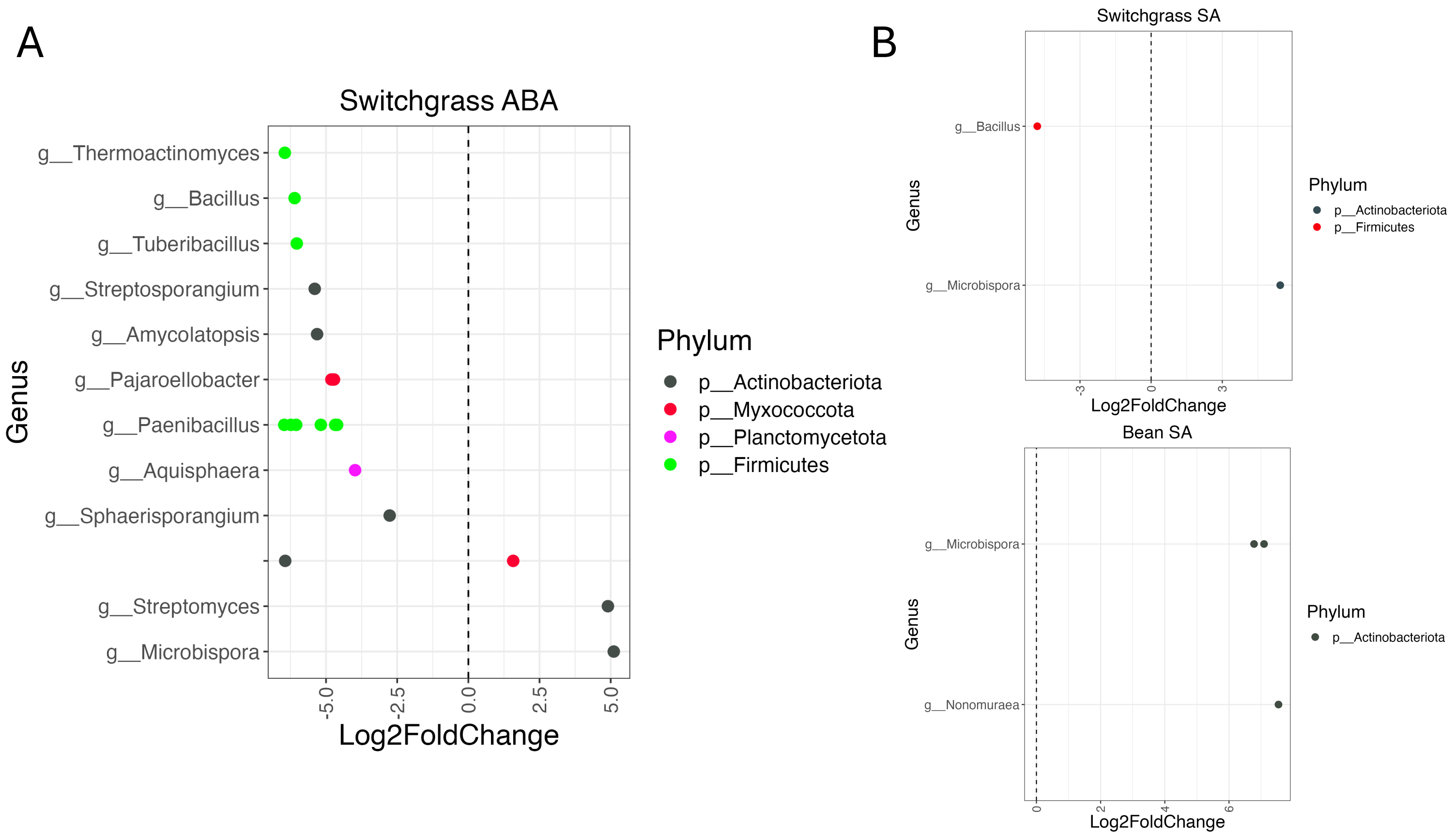


**Figure S2**. **Log two-fold enrichment and depletion of root-zone bacterial taxa that were activated in the water-acclimated versus the Day 1 post-hormone sample, as determined by the differential expression analysis DESeq2**. Points to the left of the dashed line were enriched in water-acclimated samples, and those to the right were enriched on Day 1 after the phytohormone addition. **A)** Enrichment of active taxa following the abscisic acid (ABA) addition in switchgrass root zone soil mesocosms. There were no statistically supported enrichments of active taxa in bean soil (data not shown). **B)** Enrichment of active taxa following the salicylic acid (SA) addition.

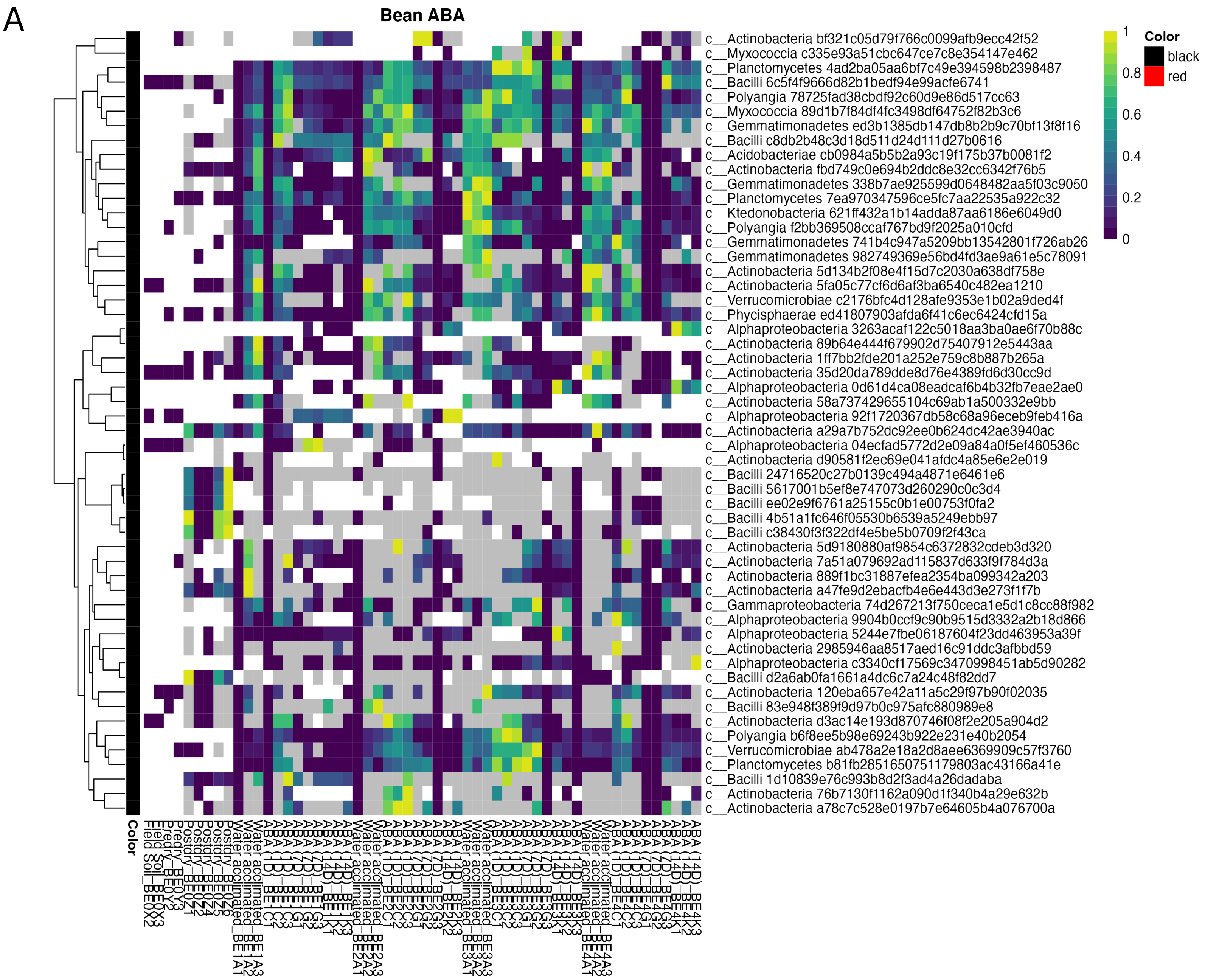
A
Bean ABA
c__Actinobacteria bf321c05d79f766c0099afb9ecc42f52
c__Myxococcia c335e93a51cbc647ce7c8e354147e462
c__Planctomycetes 4ad2ba05aa6bf7c49e394598b2398487
c__Bacilli 6c5f4f9666d82b1bedf94e99acfe6741
c__Polyangia 78725fad38cbdf92c60d9e86d517cc63
c__Myxococcia 89d1b7f84df4fc3498df64752f82b3c6
c__Gemmatimonadetes ed3b1385db147db8b2b9c70bf13f8f16
c__Bacilli c8db2b48c3d18d511d24d111d27b0616
c__Acidobacteriae cb0984a5b5b2a93c19f175b37b0081f2
c__Actinobacteria fbd749c0e694b2ddc8e32cc6342f76b5
c__Gemmatimonadetes 338b7ae925599d0648482aa5f03c9050
c__Planctomycetes 7ea970347596ce5fc7aa22535a922c32
c__Ktedonobacteria 621ff432a1b14adda87aa6186e6049d0
c__Polyangia f2bb369508ccaf767bd9f2025a010cfd
c__Gemmatimonadetes 741b4c947a5209bb13542801f726ab26
c__Gemmatimonadetes 982749369e56bd4fd3ae9a61e5c78091
c__Actinobacteria 5d134b2f08e4f15d7c2030a638df758e
c__Actinobacteria 5fa05c77cf6d6af3ba6540c482ea1210
c__Verrucomicrobiae c2176bfc4d128afe9353e1b02a9ded4f
c__Phycisphaerae ed41807903afda6f41c6ec6424cfd15a
c__Alphaproteobacteria 3263acaf122c5018aa3ba0ae6f70b88c
c__Actinobacteria 89b64e444f679902d75407912e5443aa
c__Actinobacteria 1ff7bb2fde201a252e759c8b887b265a
c__Actinobacteria 35d20da789dde8d76e4389fd6d30cc9d
c__Alphaproteobacteria 0d61d4ca08eadcaf6b4b32fb7eae2ae0
c__Actinobacteria 58a737429655104c69ab1a500332e9bb
c__Alphaproteobacteria 92f1720367db58c68a96eceb9feb416a
c__Actinobacteria a29a7b752dc92ee0b624dc42ae3940ac
c__Alphaproteobacteria 04ecfad5772d2e09a84a0f5ef460536c
c__Actinobacteria d90581f2ec69e041afdc4a85e6e2e019
c__Bacilli 24716520c27b0139c494a4871e6461e6
c__Bacilli 5617001b5ef8e747073d260290c0c3d4
c__Bacilli ee02e9f6761a25155c0b1e00753f0fa2
c__Bacilli 4b51a1fc646f05530b6539a5249ebb97
c__Bacilli c38430f3f322df4e5be5b0709f2f43ca
c__Actinobacteria 5d9180880af9854c6372832cdeb3d320
c__Actinobacteria 7a51a079692ad115837d633f9f784d3a
c__Actinobacteria 889f1bc31887efea2354ba099342a203
c__Actinobacteria a47fe9d2ebacfb4e6e443d3e273f1f7b
c__Gammaproteobacteria 74d267213f750ceca1e5d1c8cc88f982
c__Alphaproteobacteria 9904b0ccf9c90b9515d3332a2b18d866
c__Alphaproteobacteria 5244e7fbe06187604f23dd463953a39f
c__Actinobacteria 2985946aa8517aed16c91ddc3afbbd59
c__Alphaproteobacteria c3340cf17569c3470998451ab5d90282
c__Bacilli d2a6ab0fa1661a4dc6c7a24c48f82dd7
c__Actinobacteria 120eba657e42a11a5c29f97b90f02035
c__Bacilli 83e948f389f9d97b0c975afc880989e8
c__Actinobacteria d3ac14e193d870746f08f2e205a904d2
c__Polyangia b6f8ee5b98e69243b922e231e40b2054
c__Verrucomicrobiae ab478a2e18a2d8aee6369909c57f3760
c__Planctomycetes b81fb2851650751179803ac43166a41e
c__Bacilli 1d10839e76c993b8d2f3ad4a26dadaba
c__Actinobacteria 76b7130f1162a090d1f340b4a29e632b
c__Actinobacteria a78c7c528e0197b7e64605b4a076700a
Color
1
0.8
0.6
0.4
0.2
0
Color
black
red

B

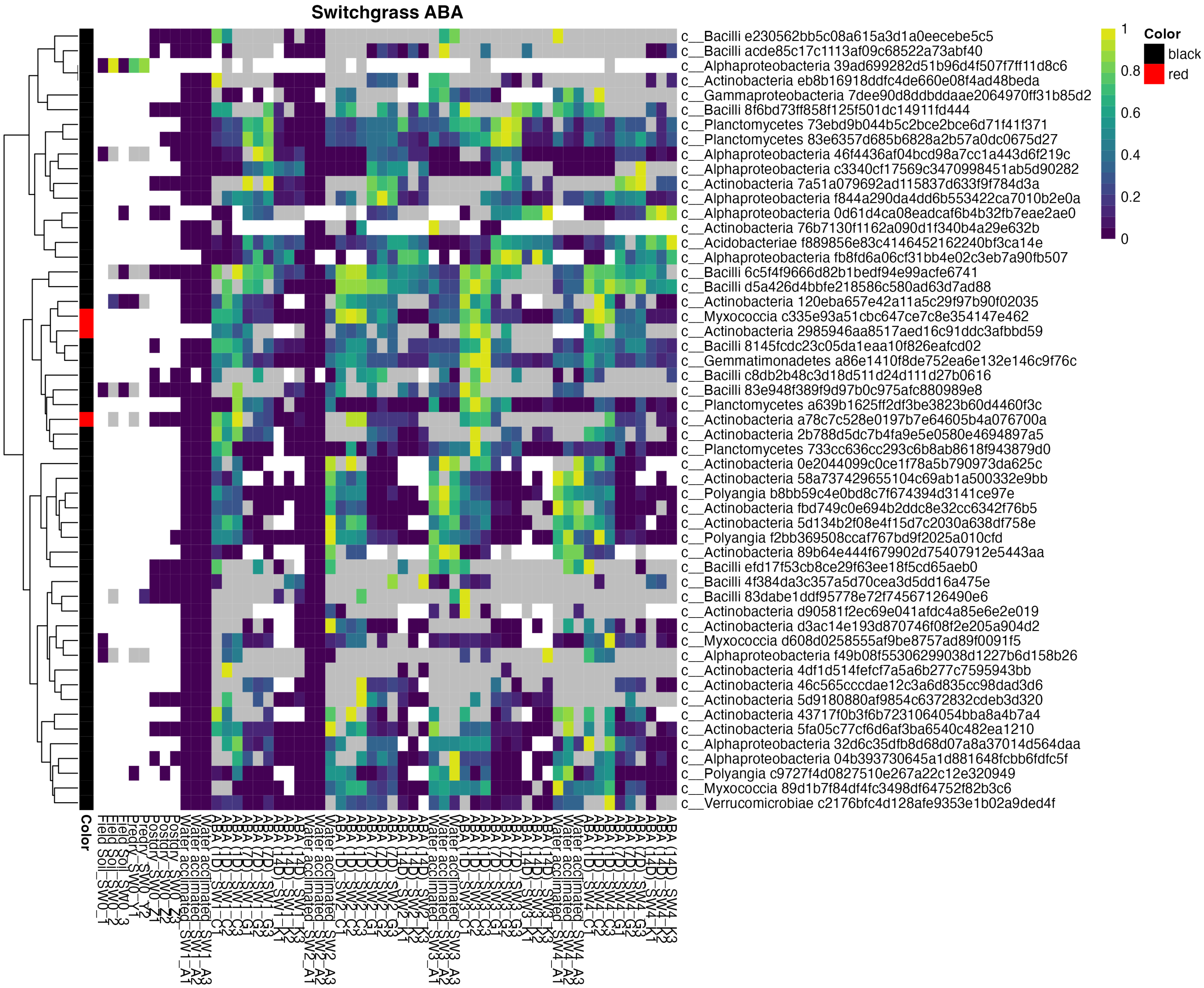

Switchgrass ABA
c__Bacilli e230562bb5c08a615a3d1a0eecebe5c5
c__Bacilli acde85c17c1113af09c68522a73abf40
c__Alphaproteobacteria 39ad699282d51b96d4f507f7ff11d8c6
c__Actinobacteria eb8b16918ddfc4de660e08f4ad48beda
c__Gammaproteobacteria 7dee90d8ddbddaae2064970ff31b85d2
c__Bacilli 8f6bd73ff858f125f501dc14911fd444
c__Planctomycetes 73ebd9b044b5c2bce2bce6d71f41f371
c__Planctomycetes 83e6357d685b6828a2b57a0dc0675d27
c__Alphaproteobacteria 46f4436af04bcd98a7cc1a443d6f219c
c__Alphaproteobacteria c3340cf17569c3470998451ab5d90282
c__Actinobacteria 7a51a079692ad115837d633f9f784d3a
c__Alphaproteobacteria f844a290da4dd6b553422ca7010b2e0a
c__Alphaproteobacteria 0d61d4ca08eadcaf6b4b32fb7eae2ae0
c__Actinobacteria 76b7130f1162a090d1f340b4a29e632b
c__Acidobacteriae f889856e83c4146452162240bf3ca14e
c__Alphaproteobacteria fb8fd6a06cf31bb4e02c3eb7a90fb507
c__Bacilli 6c5f4f9666d82b1bedf94e99acfe6741
c__Bacilli d5a426d4bbfe218586c580ad63d7ad88
c__Actinobacteria 120eba657e42a11a5c29f97b90f02035
c__Myxococcia c335e93a51cbc647ce7c8e354147e462
c__Actinobacteria 2985946aa8517aed16c91ddc3afbbd59
c__Bacilli 8145fcdc23c05da1eaa10f826eafcd02
c__Gemmatimonadetes a86e1410f8de752ea6e132e146c9f76c
c__Bacilli c8db2b48c3d18d511d24d111d27b0616
c__Bacilli 83e948f389f9d97b0c975afc880989e8
c__Planctomycetes a639b1625ff2df3be3823b60d4460f3c
c__Actinobacteria a78c7c528e0197b7e64605b4a076700a
c__Actinobacteria 2b788d5dc7b4fa9e5e0580e4694897a5
c__Planctomycetes 733cc636cc293c6b8ab8618f943879d0
c__Actinobacteria 0e2044099c0ce1f78a5b790973da625c
c__Actinobacteria 58a737429655104c69ab1a500332e9bb
c__Polyangia b8bb59c4e0bd8c7f674394d3141ce97e
c__Actinobacteria fbd749c0e694b2ddc8e32cc6342f76b5
c__Actinobacteria 5d134b2f08e4f15d7c2030a638df758e
c__Polyangia f2bb369508ccaf767bd9f2025a010cfd
c__Actinobacteria 89b64e444f679902d75407912e5443aa
c__Bacilli efd17f53cb8ce29f63ee18f5cd65aeb0
c__Bacilli 4f384da3c357a5d70cea3d5dd16a475e
c__Bacilli 83dabe1ddf95778e72f74567126490e6
c__Actinobacteria d90581f2ec69e041afdc4a85e6e2e019
c__Actinobacteria d3ac14e193d870746f08f2e205a904d2
c__Myxococcia d608d0258555af9be8757ad89f0091f5
c__Alphaproteobacteria f49b08f55306299038d1227b6d158b26
c__Actinobacteria 4df1d514fefcf7a5a6b277c7595943bb
c__Actinobacteria 46c565cccdae12c3a6d835cc98dad3d6
c__Actinobacteria 5d9180880af9854c6372832cdeb3d320
c__Actinobacteria 43717f0b3f6b7231064054bba8a4b7a4
c__Actinobacteria 5fa05c77cf6d6af3ba6540c482ea1210
c__Alphaproteobacteria 32d6c35dfb8d68d07a8a37014d564daa
c__Alphaproteobacteria 04b393730645a1d881648fcbb6fdfc5f
c__Polyangia c9727f4d0827510e267a22c12e320949
c__Myxococcia 89d1b7f84df4fc3498df64752f82b3c6
c__Verrucomicrobiae c2176bfc4d128afe9353e1b02a9ded4f
Color
1
0.8
0.6
0.4
0.2
0
Color
black
red

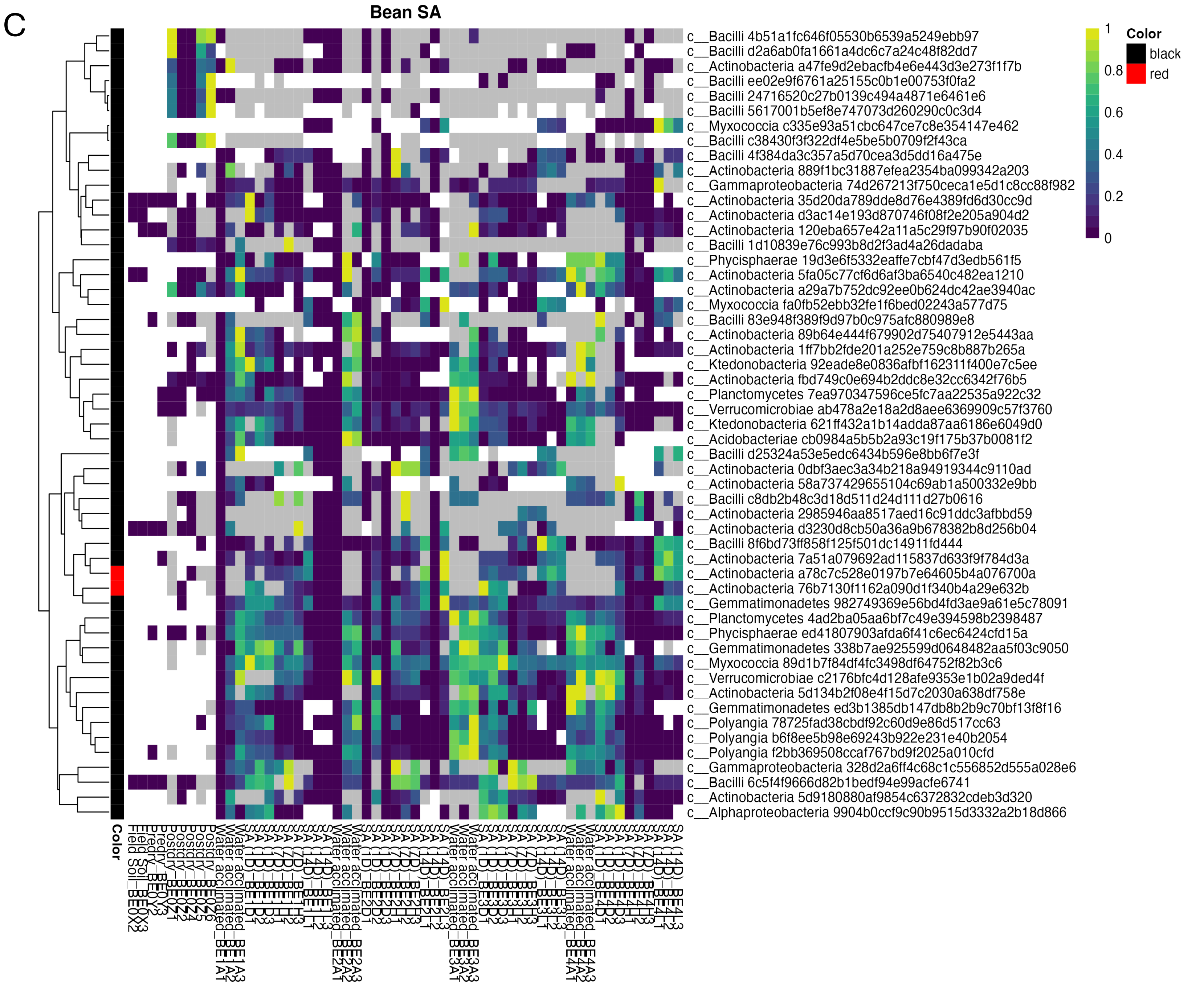

C
Bean SA
Color
black
red
1
0.8
0.6
0.4
0.2
0
c__Bacilli 4b51a1fc646f05530b6539a5249ebb97
c__Bacilli d2a6ab0fa1661a4dc6c7a24c48f82dd7
c__Actinobacteria a47fe9d2ebacfb4e6e443d3e273f1f7b
c__Bacilli ee02e9f6761a25155c0b1e00753f0fa2
c__Bacilli 24716520c27b0139c494a4871e6461e6
c__Bacilli 5617001b5ef8e747073d260290c0c3d4
c__Myxococcia c335e93a51cbc647ce7c8e354147e462
c__Bacilli c38430f3f322df4e5be5b0709f2f43ca
c__Bacilli 4f384da3c357a5d70cea3d5dd16a475e
c__Actinobacteria 889f1bc31887efea2354ba099342a203
c__Gammaproteobacteria 74d267213f750ceca1e5d1c8cc88f982
c__Actinobacteria 35d20da789dde8d76e4389fd6d30cc9d
c__Actinobacteria d3ac14e193d870746f08f2e205a904d2
c__Actinobacteria 120eba657e42a11a5c29f97b90f02035
c__Bacilli 1d10839e76c993b8d2f3ad4a26dadaba
c__Phycisphaerae 19d3e6f5332eaffe7cbf47d3edb561f5
c__Actinobacteria 5fa05c77cf6d6af3ba6540c482ea1210
c__Actinobacteria a29a7b752dc92ee0b624dc42ae3940ac
c__Myxococcia fa0fb52ebb32fe1f6bed02243a577d75
c__Bacilli 83e948f389f9d97b0c975afc880989e8
c__Actinobacteria 89b64e444f679902d75407912e5443aa
c__Actinobacteria 1ff7bb2fde201a252e759c8b887b265a
c__Ktedonobacteria 92eade8e0836afbf162311f400e7c5ee
c__Actinobacteria fbd749c0e694b2ddc8e32cc6342f76b5
c__Planctomycetes 7ea970347596ce5fc7aa22535a922c32
c__Verrucomicrobiae ab478a2e18a2d8aee6369909c57f3760
c__Ktedonobacteria 621ff432a1b14adda87aa6186e6049d0
c__Acidobacteriae cb0984a5b5b2a93c19f175b37b0081f2
c__Bacilli d25324a53e5edc6434b596e8bb6f7e3f
c__Actinobacteria 0dbf3aec3a34b218a94919344c9110ad
c__Actinobacteria 58a737429655104c69ab1a500332e9bb
c__Bacilli c8db2b48c3d18d511d24d111d27b0616
c__Actinobacteria 2985946aa8517aed16c91ddc3afbbd59
c__Actinobacteria d3230d8cb50a36a9b678382b8d256b04
c__Bacilli 8f6bd73ff858f125f501dc14911fd444
c__Actinobacteria 7a51a079692ad115837d633f9f784d3a
c__Actinobacteria a78c7c528e0197b7e64605b4a076700a
c__Actinobacteria 76b7130f1162a090d1f340b4a29e632b
c__Gemmatimonadetes 982749369e56bd4fd3ae9a61e5c78091
c__Planctomycetes 4ad2ba05aa6bf7c49e394598b2398487
c__Phycisphaerae ed41807903afda6f41c6ec6424cfd15a
c__Gemmatimonadetes 338b7ae925599d0648482aa5f03c9050
c__Myxococcia 89d1b7f84df4fc3498df64752f82b3c6
c__Verrucomicrobiae c2176bfc4d128afe9353e1b02a9ded4f
c__Actinobacteria 5d134b2f08e4f15d7c2030a638df758e
c__Gemmatimonadetes ed3b1385db147db8b2b9c70bf13f8f16
c__Polyangia 78725fad38cbdf92c60d9e86d517cc63
c__Polyangia b6f8ee5b98e69243b922e231e40b2054
c__Polyangia f2bb369508ccaf767bd9f2025a010cfd
c__Gammaproteobacteria 328d2a6ff4c68c1c556852d555a028e6
c__Bacilli 6c5f4f9666d82b1bedf94e99acfe6741
c__Actinobacteria 5d9180880af9854c6372832cdeb3d320
c__Alphaproteobacteria 9904b0ccf9c90b9515d3332a2b18d866

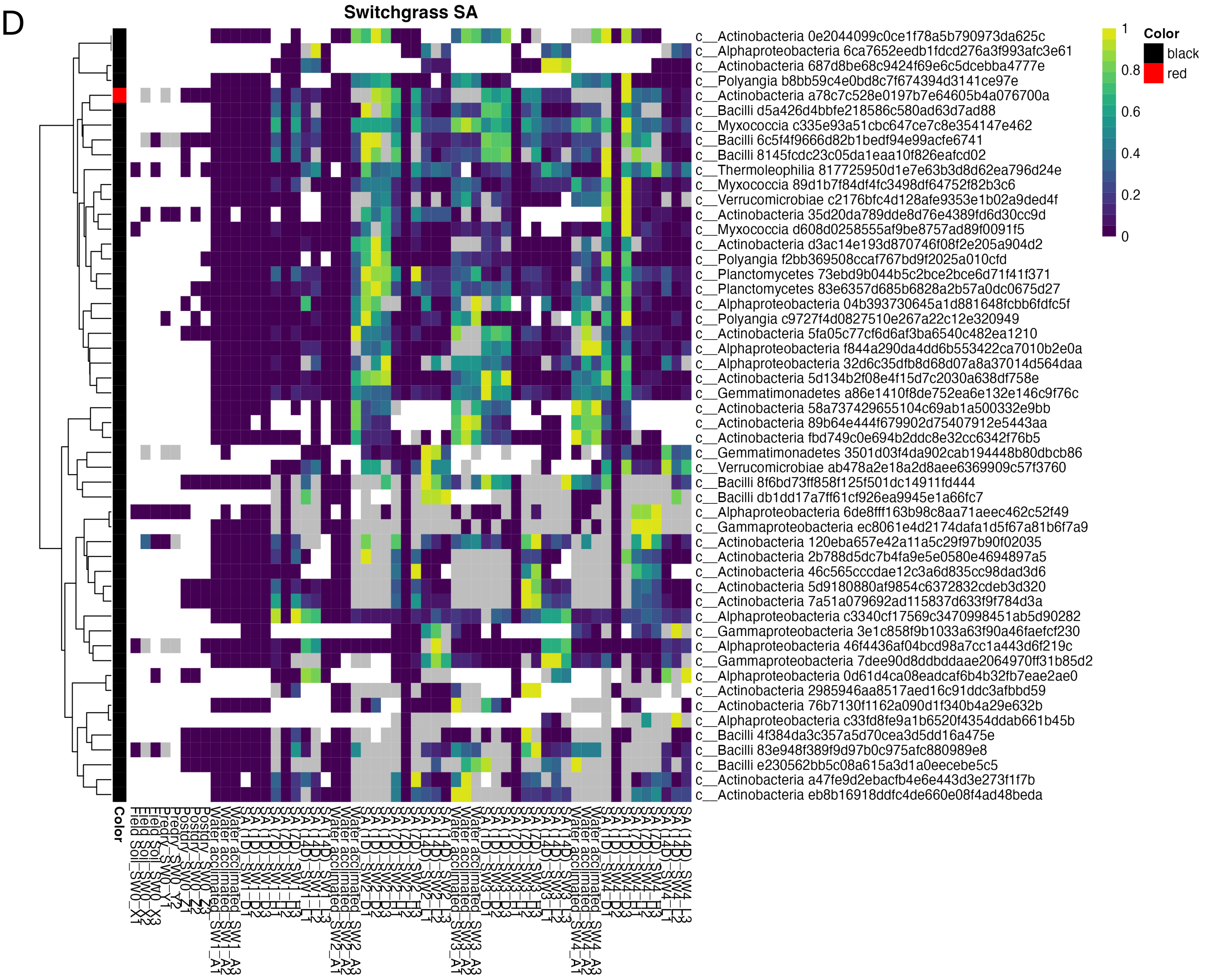
D
Switchgrass SA
Color
black
red
1
0.8
0.6
0.4
0.2
0
c__Actinobacteria 0e2044099c0ce1f78a5b790973da625c
c__Alphaproteobacteria 6ca7652eedb1fdcd276a3f993afc3e61
c__Actinobacteria 687d8be68c9424f69e6c5dcebba4777e
c__Polyangia b8bb59c4e0bd8c7f674394d3141ce97e
c__Actinobacteria a78c7c528e0197b7e64605b4a076700a
c__Bacilli d5a426d4bbfe218586c580ad63d7ad88
c__Myxococcia c335e93a51cbc647ce7c8e354147e462
c__Bacilli 6c5f4f9666d82b1bedf94e99acfe6741
c__Bacilli 8145fcdc23c05da1eaa10f826eafcd02
c__Thermoleophilia 817725950d1e7e63b3d8d62ea796d24e
c__Myxococcia 89d1b7f84df4fc3498df64752f82b3c6
c__Verrucomicrobiae c2176bfc4d128afe9353e1b02a9ded4f
c__Actinobacteria 35d20da789dde8d76e4389fd6d30cc9d
c__Myxococcia d608d0258555af9be8757ad89f0091f5
c__Actinobacteria d3ac14e193d870746f08f2e205a904d2
c__Polyangia f2bb369508ccaf767bd9f2025a010cfd
c__Planctomycetes 73ebd9b044b5c2bce2bce6d71f41f371
c__Planctomycetes 83e6357d685b6828a2b57a0dc0675d27
c__Alphaproteobacteria 04b393730645a1d881648fcbb6fdfc5f
c__Polyangia c9727f4d0827510e267a22c12e320949
c__Actinobacteria 5fa05c77cf6d6af3ba6540c482ea1210
c__Alphaproteobacteria f844a290da4dd6b553422ca7010b2e0a
c__Alphaproteobacteria 32d6c35dfb8d68d07a8a37014d564daa
c__Actinobacteria 5d134b2f08e4f15d7c2030a638df758e
c__Gemmatimonadetes a86e1410f8de752ea6e132e146c9f76c
c__Actinobacteria 58a737429655104c69ab1a500332e9bb
c__Actinobacteria 89b64e444f679902d75407912e5443aa
c__Actinobacteria fbd749c0e694b2ddc8e32cc6342f76b5
c__Gemmatimonadetes 3501d03f4da902cab194448b80dbcb86
c__Verrucomicrobiae ab478a2e18a2d8aee6369909c57f3760
c__Bacilli 8f6bd73ff858f125f501dc14911fd444
c__Bacilli db1dd17a7ff61cf926ea9945e1a66fc7
c__Alphaproteobacteria 6de8fff163b98c8aa71aeec462c52f49
c__Gammaproteobacteria ec8061e4d2174dafa1d5f67a81b6f7a9
c__Actinobacteria 120eba657e42a11a5c29f97b90f02035
c__Actinobacteria 2b788d5dc7b4fa9e5e0580e4694897a5
c__Actinobacteria 46c565cccdae12c3a6d835cc98dad3d6
c__Actinobacteria 5d9180880af9854c6372832cdeb3d320
c__Actinobacteria 7a51a079692ad115837d633f9f784d3a
c__Alphaproteobacteria c3340cf17569c3470998451ab5d90282
c__Gammaproteobacteria 3e1c858f9b1033a63f90a46faefcf230
c__Alphaproteobacteria 46f4436af04bcd98a7cc1a443d6f219c
c__Gammaproteobacteria 7dee90d8ddbddaae2064970ff31b85d2
c__Alphaproteobacteria 0d61d4ca08eadcaf6b4b32fb7eae2ae0
c__Actinobacteria 2985946aa8517aed16c91ddc3afbbd59
c__Actinobacteria 76b7130f1162a090d1f340b4a29e632b
c__Alphaproteobacteria c33fd8fe9a1b6520f4354ddab661b45b
c__Bacilli 4f384da3c357a5d70cea3d5dd16a475e
c__Bacilli 83e948f389f9d97b0c975afc880989e8
c__Bacilli e230562bb5c08a615a3d1a0eecebe5c5
c__Actinobacteria a47fe9d2ebacfb4e6e443d3e273f1f7b
c__Actinobacteria eb8b16918ddfc4de660e08f4ad48beda
Color

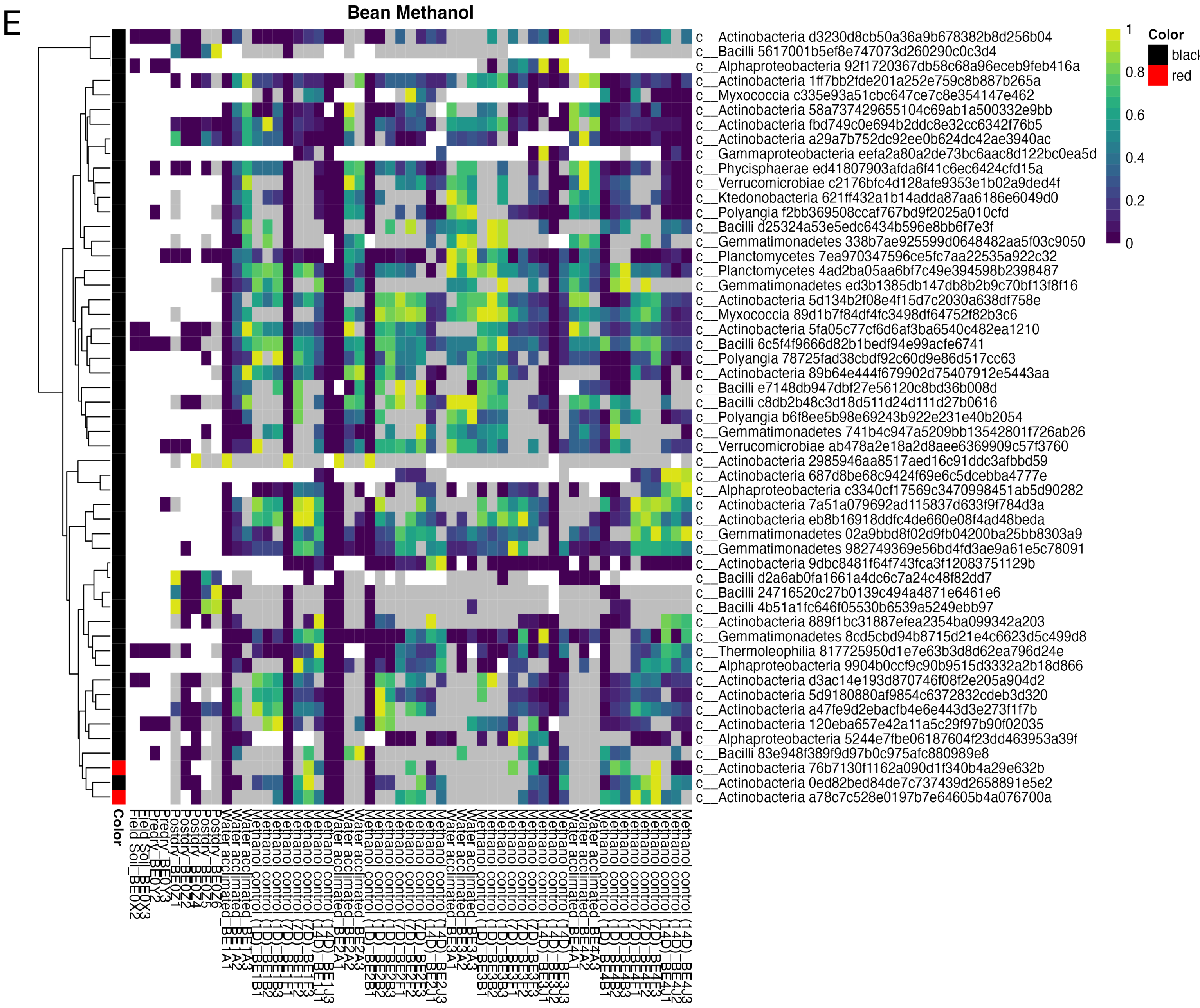
E
Bean Methanol
Color
black
red
1
0.8
0.6
0.4
0.2
0
c__Actinobacteria d3230d8cb50a36a9b678382b8d256b04
c__Bacilli 5617001b5ef8e747073d260290c0c3d4
c__Alphaproteobacteria 92f1720367db58c68a96eceb9feb416a
c__Actinobacteria 1ff7bb2fde201a252e759c8b887b265a
c__Myxococcia c335e93a51cbc647ce7c8e354147e462
c__Actinobacteria 58a737429655104c69ab1a500332e9bb
c__Actinobacteria fbd749c0e694b2ddc8e32cc6342f76b5
c__Actinobacteria a29a7b752dc92ee0b624dc42ae3940ac
c__Gammaproteobacteria eefa2a80a2de73bc6aac8d122bc0ea5d
c__Phycisphaerae ed41807903afda6f41c6ec6424cfd15a
c__Verrucomicrobiae c2176bfc4d128afe9353e1b02a9ded4f
c__Ktedonobacteria 621ff432a1b14adda87aa6186e6049d0
c__Polyangia f2bb369508ccaf767bd9f2025a010cfd
c__Bacilli d25324a53e5edc6434b596e8bb6f7e3f
c__Gemmatimonadetes 338b7ae925599d0648482aa5f03c9050
c__Planctomycetes 7ea970347596ce5fc7aa22535a922c32
c__Planctomycetes 4ad2ba05aa6bf7c49e394598b2398487
c__Gemmatimonadetes ed3b1385db147db8b2b9c70bf13f8f16
c__Actinobacteria 5d134b2f08e4f15d7c2030a638df758e
c__Myxococcia 89d1b7f84df4fc3498df64752f82b3c6
c__Actinobacteria 5fa05c77cf6d6af3ba6540c482ea1210
c__Bacilli 6c5f4f9666d82b1bedf94e99acfe6741
c__Polyangia 78725fad38cbdf92c60d9e86d517cc63
c__Actinobacteria 89b64e444f679902d75407912e5443aa
c__Bacilli e7148db947dbf27e56120c8bd36b008d
c__Bacilli c8db2b48c3d18d511d24d111d27b0616
c__Polyangia b6f8ee5b98e69243b922e231e40b2054
c__Gemmatimonadetes 741b4c947a5209bb13542801f726ab26
c__Verrucomicrobiae ab478a2e18a2d8aee6369909c57f3760
c__Actinobacteria 2985946aa8517aed16c91ddc3afbbd59
c__Actinobacteria 687d8be68c9424f69e6c5dcebba4777e
c__Alphaproteobacteria c3340cf17569c3470998451ab5d90282
c__Actinobacteria 7a51a079692ad115837d633f9f784d3a
c__Actinobacteria eb8b16918ddfc4de660e08f4ad48beda
c__Gemmatimonadetes 02a9bbd8f02d9fb04200ba25bb8303a9
c__Gemmatimonadetes 982749369e56bd4fd3ae9a61e5c78091
c__Actinobacteria 9dbc8481f64f743fca3f12083751129b
c__Bacilli d2a6ab0fa1661a4dc6c7a24c48f82dd7
c__Bacilli 24716520c27b0139c494a4871e6461e6
c__Bacilli 4b51a1fc646f05530b6539a5249ebb97
c__Actinobacteria 889f1bc31887efea2354ba099342a203
c__Gemmatimonadetes 8cd5cbd94b8715d21e4c6623d5c499d8
c__Thermoleophilia 817725950d1e7e63b3d8d62ea796d24e
c__Alphaproteobacteria 9904b0ccf9c90b9515d3332a2b18d866
c__Actinobacteria d3ac14e193d870746f08f2e205a904d2
c__Actinobacteria 5d9180880af9854c6372832cdeb3d320
c__Actinobacteria a47fe9d2ebacfb4e6e443d3e273f1f7b
c__Actinobacteria 120eba657e42a11a5c29f97b90f02035
c__Alphaproteobacteria 5244e7fbe06187604f23dd463953a39f
c__Bacilli 83e948f389f9d97b0c975afc880989e8
c__Actinobacteria 76b7130f1162a090d1f340b4a29e632b
c__Actinobacteria 0ed82bed84de7c737439d2658891e5e2
c__Actinobacteria a78c7c528e0197b7e64605b4a076700a
Color

F

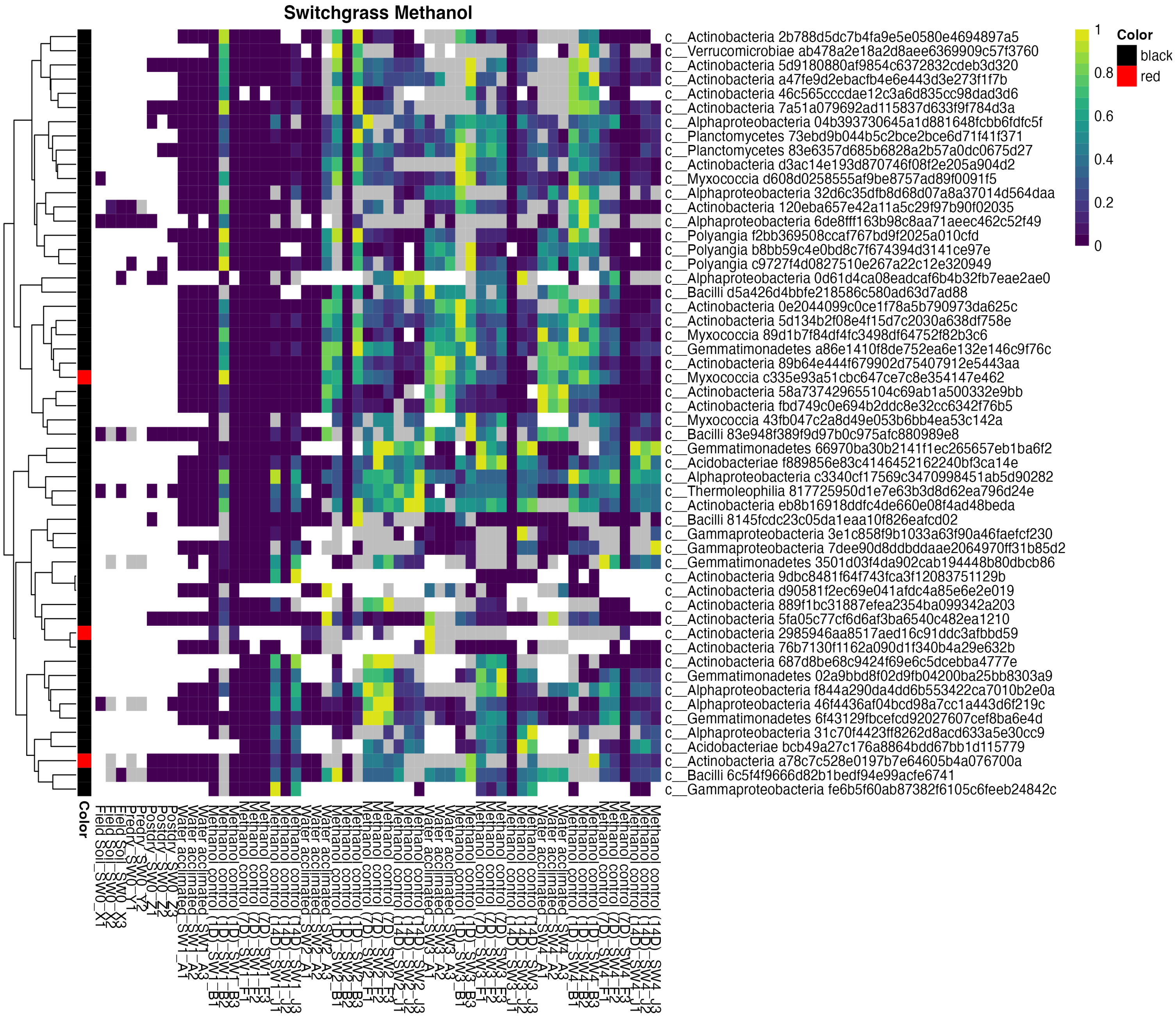
Switchgrass Methanol
c__Actinobacteria 2b788d5dc7b4fa9e5e0580e4694897a5
c__Verrucomicrobiae ab478a2e18a2d8aee6369909c57f3760
c__Actinobacteria 5d9180880af9854c6372832cdeb3d320
c__Actinobacteria a47fe9d2ebacfb4e6e443d3e273f1f7b
c__Actinobacteria 46c565cccdae12c3a6d835cc98dad3d6
c__Actinobacteria 7a51a079692ad115837d633f9f784d3a
c__Alphaproteobacteria 04b393730645a1d881648fcbb6fdfc5f
c__Planctomycetes 73ebd9b044b5c2bce2bce6d71f41f371
c__Planctomycetes 83e6357d685b6828a2b57a0dc0675d27
c__Actinobacteria d3ac14e193d870746f08f2e205a904d2
c__Myxococcia d608d0258555af9be8757ad89f0091f5
c__Alphaproteobacteria 32d6c35dfb8d68d07a8a37014d564daa
c__Actinobacteria 120eba657e42a11a5c29f97b90f02035
c__Alphaproteobacteria 6de8fff163b98c8aa71aeec462c52f49
c__Polyangia f2bb369508ccaf767bd9f2025a010cfd
c__Polyangia b8bb59c4e0bd8c7f674394d3141ce97e
c__Polyangia c9727f4d0827510e267a22c12e320949
c__Alphaproteobacteria 0d61d4ca08eadcaf6b4b32fb7eae2ae0
c__Bacilli d5a426d4bbfe218586c580ad63d7ad88
c__Actinobacteria 0e2044099c0ce1f78a5b790973da625c
c__Actinobacteria 5d134b2f08e4f15d7c2030a638df758e
c__Myxococcia 89d1b7f84df4fc3498df64752f82b3c6
c__Gemmatimonadetes a86e1410f8de752ea6e132e146c9f76c
c__Actinobacteria 89b64e444f679902d75407912e5443aa
c__Myxococcia c335e93a51cbc647ce7c8e354147e462
c__Actinobacteria 58a737429655104c69ab1a500332e9bb
c__Actinobacteria fbd749c0e694b2ddc8e32cc6342f76b5
c__Myxococcia 43fb047c2a8d49e053b6bb4ea53c142a
c__Bacilli 83e948f389f9d97b0c975afc880989e8
c__Gemmatimonadetes 66970ba30b2141f1ec265657eb1ba6f2
c__Acidobacteriae f889856e83c4146452162240bf3ca14e
c__Alphaproteobacteria c3340cf17569c3470998451ab5d90282
c__Thermoleophilia 817725950d1e7e63b3d8d62ea796d24e
c__Actinobacteria eb8b16918ddfc4de660e08f4ad48beda
c__Bacilli 8145fcdc23c05da1eaa10f826eafcd02
c__Gammaproteobacteria 3e1c858f9b1033a63f90a46faefcf230
c__Gammaproteobacteria 7dee90d8ddbddaae2064970ff31b85d2
c__Gemmatimonadetes 3501d03f4da902cab194448b80dbcb86
c__Actinobacteria 9dbc8481f64f743fca3f12083751129b
c__Actinobacteria d90581f2ec69e041afdc4a85e6e2e019
c__Actinobacteria 889f1bc31887efea2354ba099342a203
c__Actinobacteria 5fa05c77cf6d6af3ba6540c482ea1210
c__Actinobacteria 2985946aa8517aed16c91ddc3afbbd59
c__Actinobacteria 76b7130f1162a090d1f340b4a29e632b
c__Actinobacteria 687d8be68c9424f69e6c5dcebba4777e
c__Gemmatimonadetes 02a9bbd8f02d9fb04200ba25bb8303a9
c__Alphaproteobacteria f844a290da4dd6b553422ca7010b2e0a
c__Alphaproteobacteria 46f4436af04bcd98a7cc1a443d6f219c
c__Gemmatimonadetes 6f43129fbcefcd92027607cef8ba6e4d
c__Alphaproteobacteria 31c70f4423ff8262d8acd633a5e30cc9
c__Acidobacteriae bcb49a27c176a8864bdd67bb1d115779
c__Actinobacteria a78c7c528e0197b7e64605b4a076700a
c__Bacilli 6c5f4f9666d82b1bedf94e99acfe6741
c__Gammaproteobacteria fe6b5f60ab87382f6105c6feeb24842c
1
0.8
0.6
0.4
0.2
0
Color
black
red
Color

**Figure S3. Activity dynamics of the most abundant 50 taxa observed for the bean and switchgrass root zone soils exposed to phytohormone treatments (abscisic acid, ABA, and salicylic acid, SA) and methanol carrier controls. All mesocosms are shown separately for A) Bean ABA, B) Switchgrass ABA, C) Bean SA, D) Switchgrass SA, E) Bean methanol, F) Switchgrass methanol**. Relative abundance of active members is shown with a blue (low relative abundance) to yellow (high relative abundance) color gradient. Inactive taxa detected in DNA dataset (DNA>cDNA) are denoted with grey and corresponds to a zero abundance. Members not detected in DNA or cDNA count tables are colored as white and also denote zero active abundance. Red and black markers along the rows indicate which taxa were part of the log two-fold enriched group (enriched in Day 1 compared to water-acclimated samples after phytohormone addition) by DESeq analysis, red showing enrichment, black showing no enrichment. If these taxa were not in the top 50 abundant pool, they were separately added for comparisons.

A

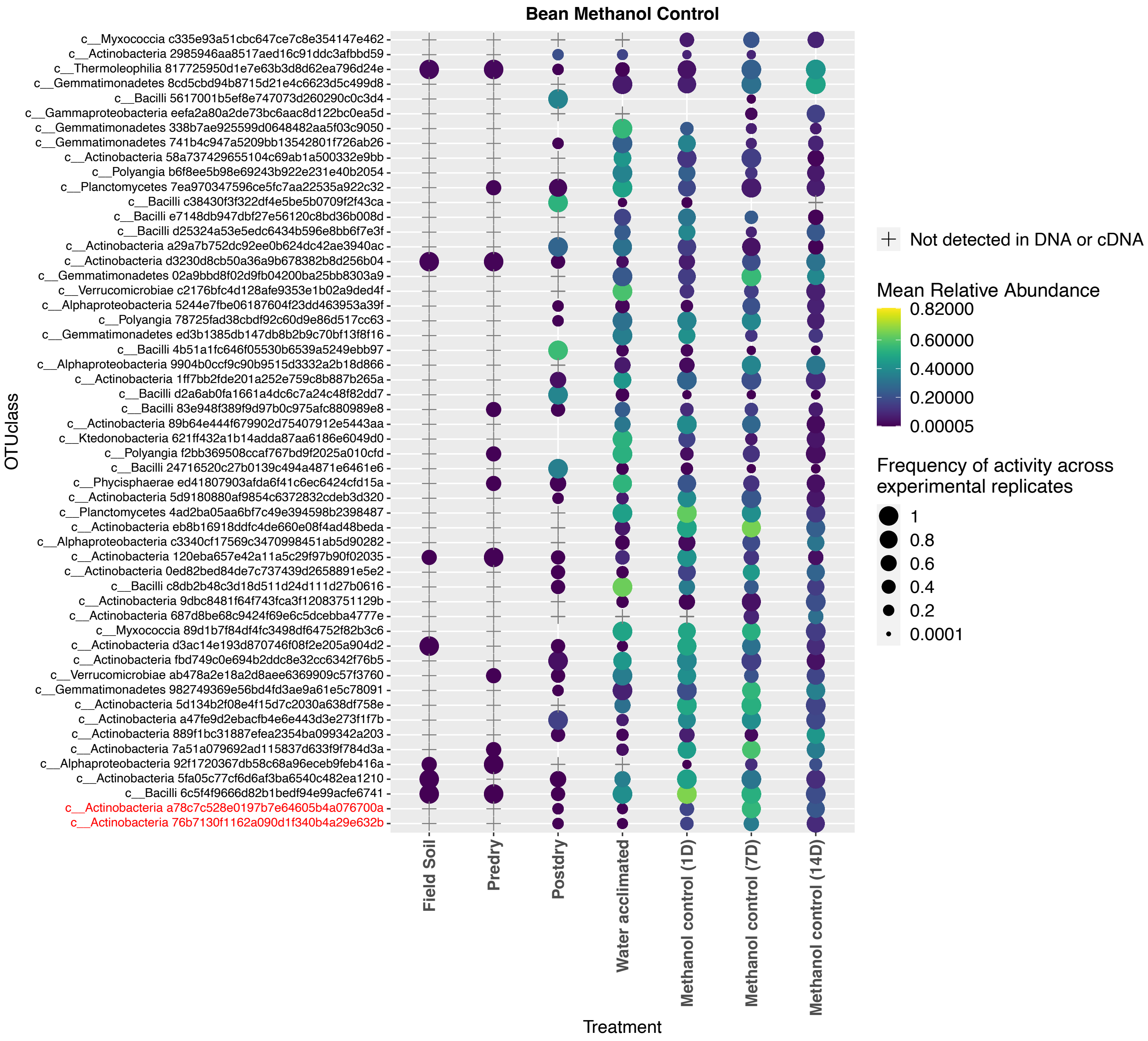

Bean Methanol Control
OTUclass
c__Myxococcia c335e93a51cbc647ce7c8e354147e462
c__Actinobacteria 2985946aa8517aed16c91ddc3afbbd59
c__Thermoleophilia 817725950d1e7e63b3d8d62ea796d24e
c__Gemmatimonadetes 8cd5cbd94b8715d21e4c6623d5c499d8
c__Bacilli 5617001b5ef8e747073d260290c0c3d4
c__Gammaproteobacteria eefa2a80a2de73bc6aac8d122bc0ea5d
c__Gemmatimonadetes 338b7ae925599d0648482aa5f03c9050
c__Gemmatimonadetes 741b4c947a5209bb13542801f726ab26
c__Actinobacteria 58a737429655104c69ab1a500332e9bb
c__Polyangia b6f8ee5b98e69243b922e231e40b2054
c__Planctomycetes 7ea970347596ce5fc7aa22535a922c32
c__Bacilli c38430f3f322df4e5be5b0709f2f43ca
c__Bacilli e7148db947dbf27e56120c8bd36b008d
c__Bacilli d25324a53e5edc6434b596e8bb6f7e3f
c__Actinobacteria a29a7b752dc92ee0b624dc42ae3940ac
c__Actinobacteria d3230d8cb50a36a9b678382b8d256b04
c__Gemmatimonadetes 02a9bbd8f02d9fb04200ba25bb8303a9
c__Verrucomicrobiae c2176bfc4d128afe9353e1b02a9ded4f
c__Alphaproteobacteria 5244e7fbe06187604f23dd463953a39f
c__Polyangia 78725fad38cbdf92c60d9e86d517cc63
c__Gemmatimonadetes ed3b1385db147db8b2b9c70bf13f8f16
c__Bacilli 4b51a1fc646f05530b6539a5249ebb97
c__Alphaproteobacteria 9904b0ccf9c90b9515d3332a2b18d866
c__Actinobacteria 1ff7bb2fde201a252e759c8b887b265a
c__Bacilli d2a6ab0fa1661a4dc6c7a24c48f82dd7
c__Bacilli 83e948f389f9d97b0c975afc880989e8
c__Actinobacteria 89b64e444f679902d75407912e5443aa
c__Ktedonobacteria 621ff432a1b14adda87aa6186e6049d0
c__Polyangia f2bb369508ccaf767bd9f2025a010cfd
c__Bacilli 24716520c27b0139c494a4871e6461e6
c__Phycisphaerae ed41807903afda6f41c6ec6424cfd15a
c__Actinobacteria 5d9180880af9854c6372832cdeb3d320
c__Planctomycetes 4ad2ba05aa6bf7c49e394598b2398487
c__Actinobacteria eb8b16918ddfc4de660e08f4ad48beda
c__Alphaproteobacteria c3340cf17569c3470998451ab5d90282
c__Actinobacteria 120eba657e42a11a5c29f97b90f02035
c__Actinobacteria 0ed82bed84de7c737439d2658891e5e2
c__Bacilli c8db2b48c3d18d511d24d111d27b0616
c__Actinobacteria 9dbc8481f64f743fca3f12083751129b
c__Actinobacteria 687d8be68c9424f69e6c5dcebba4777e
c__Myxococcia 89d1b7f84df4fc3498df64752f82b3c6
c__Actinobacteria d3ac14e193d870746f08f2e205a904d2
c__Actinobacteria fbd749c0e694b2ddc8e32cc6342f76b5
c__Verrucomicrobiae ab478a2e18a2d8aee6369909c57f3760
c__Gemmatimonadetes 982749369e56bd4fd3ae9a61e5c78091
c__Actinobacteria 5d134b2f08e4f15d7c2030a638df758e
c__Actinobacteria a47fe9d2ebacfb4e6e443d3e273f1f7b
c__Actinobacteria 889f1bc31887efea2354ba099342a203
c__Actinobacteria 7a51a079692ad115837d633f9f784d3a
c__Alphaproteobacteria 92f1720367db58c68a96eceb9feb416a
c__Actinobacteria 5fa05c77cf6d6af3ba6540c482ea1210
c__Bacilli 6c5f4f9666d82b1bedf94e99acfe6741
c__Actinobacteria a78c7c528e0197b7e64605b4a076700a
c__Actinobacteria 76b7130f1162a090d1f340b4a29e632b
Field Soil
Predry
Postdry
Water acclimated
Methanol control (1D)
Methanol control (7D)
Methanol control (14D)
Treatment
Not detected in DNA or cDNA
Mean Relative Abundance
0.82000
0.60000
0.40000
0.20000
0.00005
Frequency of activity across experimental replicates
1
0.8
0.6
0.4
0.2
0.0001

B

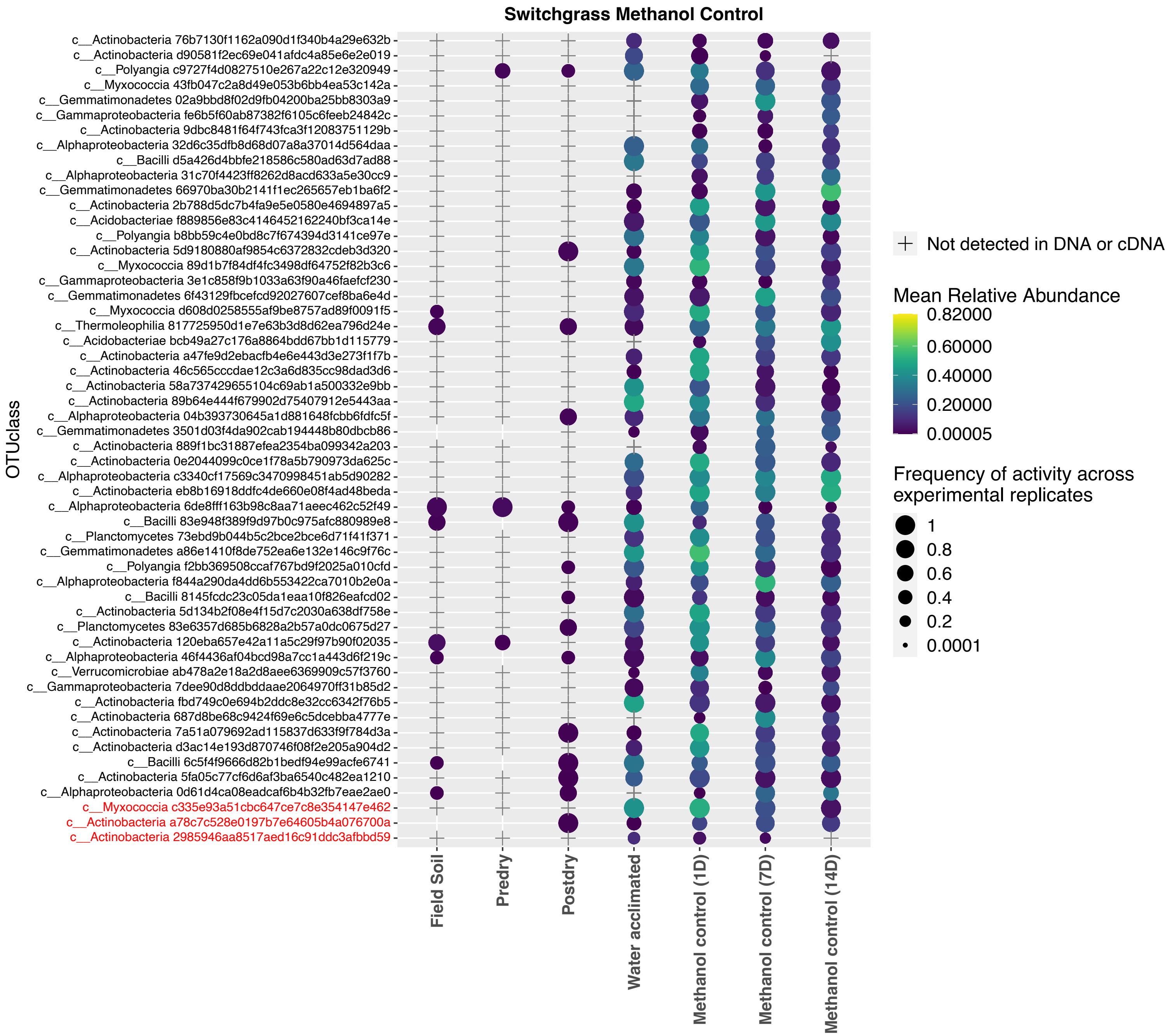
Switchgrass Methanol Control
c__Actinobacteria 76b7130f1162a090d1f340b4a29e632b
c__Actinobacteria d90581f2ec69e041afdc4a85e6e2e019
c__Polyangia c9727f4d0827510e267a22c12e320949
c__Myxococcia 43fb047c2a8d49e053b6bb4ea53c142a
c__Gemmatimonadetes 02a9bbd8f02d9fb04200ba25bb8303a9
c__Gammaproteobacteria fe6b5f60ab87382f6105c6feeb24842c
c__Actinobacteria 9dbc8481f64f743fca3f12083751129b
c__Alphaproteobacteria 32d6c35dfb8d68d07a8a37014d564daa
c__Bacilli d5a426d4bbfe218586c580ad63d7ad88
c__Alphaproteobacteria 31c70f4423ff8262d8acd633a5e30cc9
c__Gemmatimonadetes 66970ba30b2141f1ec265657eb1ba6f2
c__Actinobacteria 2b788d5dc7b4fa9e5e0580e4694897a5
c__Acidobacteriae f889856e83c4146452162240bf3ca14e
c__Polyangia b8bb59c4e0bd8c7f674394d3141ce97e
c__Actinobacteria 5d9180880af9854c6372832cdeb3d320
c__Myxococcia 89d1b7f84df4fc3498df64752f82b3c6
c__Gammaproteobacteria 3e1c858f9b1033a63f90a46faefcf230
c__Gemmatimonadetes 6f43129fbcefcd92027607cef8ba6e4d
c__Myxococcia d608d0258555af9be8757ad89f0091f5
c__Thermoleophilia 817725950d1e7e63b3d8d62ea796d24e
c__Acidobacteriae bcb49a27c176a8864bdd67bb1d115779
c__Actinobacteria a47fe9d2ebacfb4e6e443d3e273f1f7b
c__Actinobacteria 46c565cccdae12c3a6d835cc98dad3d6
c__Actinobacteria 58a737429655104c69ab1a500332e9bb
c__Actinobacteria 89b64e444f679902d75407912e5443aa
c__Alphaproteobacteria 04b393730645a1d881648fcbb6fdfc5f
c__Gemmatimonadetes 3501d03f4da902cab194448b80dbcb86
c__Actinobacteria 889f1bc31887efea2354ba099342a203
c__Actinobacteria 0e2044099c0ce1f78a5b790973da625c
c__Alphaproteobacteria c3340cf17569c3470998451ab5d90282
c__Actinobacteria eb8b16918ddfc4de660e08f4ad48beda
c__Alphaproteobacteria 6de8fff163b98c8aa71aeec462c52f49
c__Bacilli 83e948f389f9d97b0c975afc880989e8
c__Planctomycetes 73ebd9b044b5c2bce2bce6d71f41f371
c__Gemmatimonadetes a86e1410f8de752ea6e132e146c9f76c
c__Polyangia f2bb369508ccaf767bd9f2025a010cfd
c__Alphaproteobacteria f844a290da4dd6b553422ca7010b2e0a
c__Bacilli 8145fcdc23c05da1eaa10f826eafcd02
c__Actinobacteria 5d134b2f08e4f15d7c2030a638df758e
c__Planctomycetes 83e6357d685b6828a2b57a0dc0675d27
c__Actinobacteria 120eba657e42a11a5c29f97b90f02035
c__Alphaproteobacteria 46f4436af04bcd98a7cc1a443d6f219c
c__Verrucomicrobiae ab478a2e18a2d8aee6369909c57f3760
c__Gammaproteobacteria 7dee90d8ddbddaae2064970ff31b85d2
c__Actinobacteria fbd749c0e694b2ddc8e32cc6342f76b5
c__Actinobacteria 687d8be68c9424f69e6c5dcebba4777e
c__Actinobacteria 7a51a079692ad115837d633f9f784d3a
c__Actinobacteria d3ac14e193d870746f08f2e205a904d2
c__Bacilli 6c5f4f9666d82b1bedf94e99acfe6741
c__Actinobacteria 5fa05c77cf6d6af3ba6540c482ea1210
c__Alphaproteobacteria 0d61d4ca08eadcaf6b4b32fb7eae2ae0
c__Myxococcia c335e93a51cbc647ce7c8e354147e462
c__Actinobacteria a78c7c528e0197b7e64605b4a076700a
c__Actinobacteria 2985946aa8517aed16c91ddc3afbbd59
OTUclass
Field Soil
Predry
Postdry
Water acclimated
Methanol control (1D)
Methanol control (7D)
Methanol control (14D)
Treatment
Not detected in DNA or cDNA
Mean Relative Abundance
0.82000
0.60000
0.40000
0.20000
0.00005
Frequency of activity across experimental replicates
1
0.8
0.6
0.4
0.2
0.0001

**Figure S4. Shifts in microbial activation before and after the carrier control methanol addition in A) bean and B) switchgrass soil mesocosms with responses aggregated across replicate mesocosms**. Relative abundances of active members that were enriched at Day 1 after phytohormone addition compared to the water-acclimated sample are shown in red text and were part of the log two-fold enriched group by DESeq analysis. All other taxa were the 50 most abundant active members in the specific treatment communities. The differences in activity between the water-acclimated and Day 1 timepoint as plotted here are specific to a methanol exposure and the red colored taxa are plotted to compare methanol response to phytohormone responses (Figure 2 in main manuscript). These comparisons indicate that these taxa were not different in activity between water-acclimated and Day 1 timepoints in methanol carrier controls but were enriched in the phytohormone treatments. Inactive taxa have a relative abundance of zero, so they do not contain a symbol on the graph and are left blank for that time point/treatment. Taxa not detected in either DNA or cDNA data set are denoted by a + shape. Color gradient denotes increased relative abundances as color changes from blue to yellow. The shape size of each bubble denotes the consistency of detection across experimental and technical replicates of soil mesocosms (e.g. with 1 denoting activity in all replicates and 0.5 denoting activity in half). Empty spaces in rows that also have symbols indicate that the taxon was not classified as active within those samples (columns) but was detected in DNA.

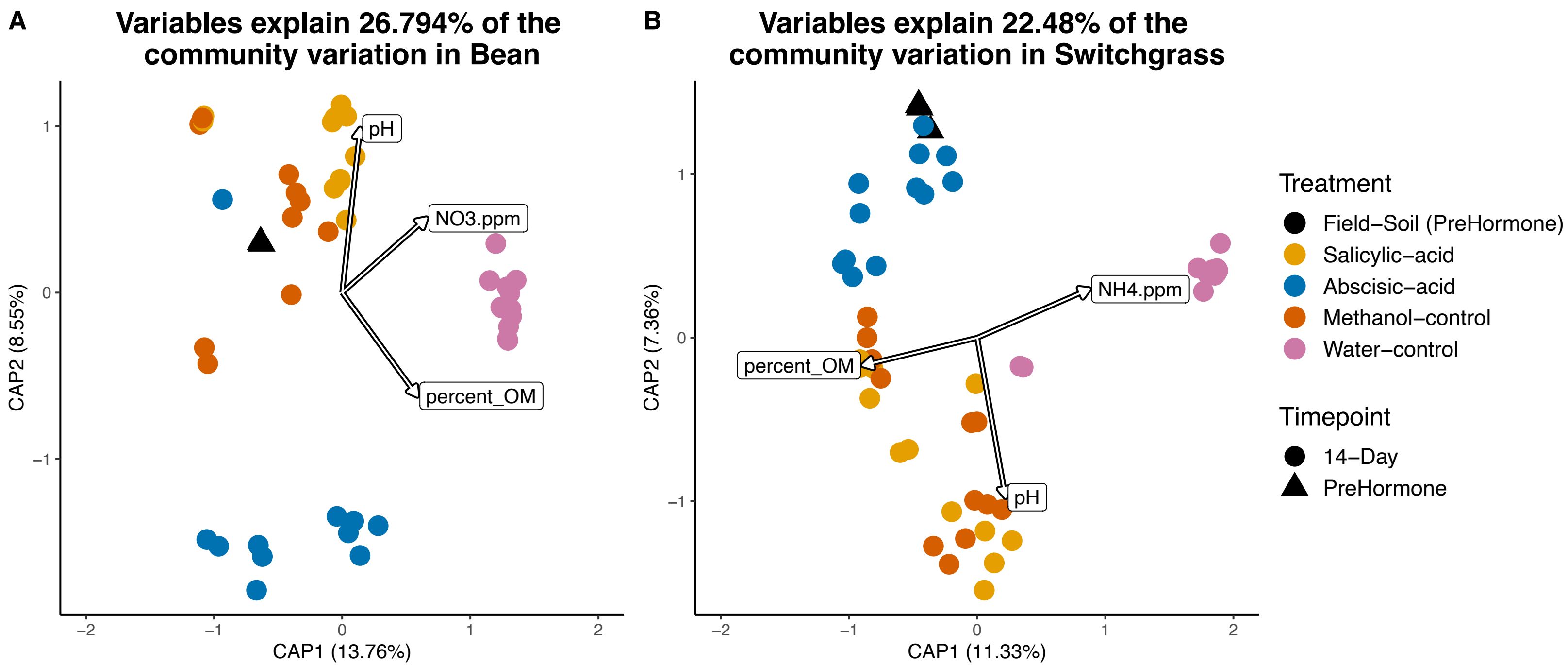


**Figure S5.** Constrained analysis of principal coordinates showing effect of soil pH, nitrate, ammonium and organic matter content on soil microbial communities (at 14-day time point) in A) Bean, and B) Switchgrass.

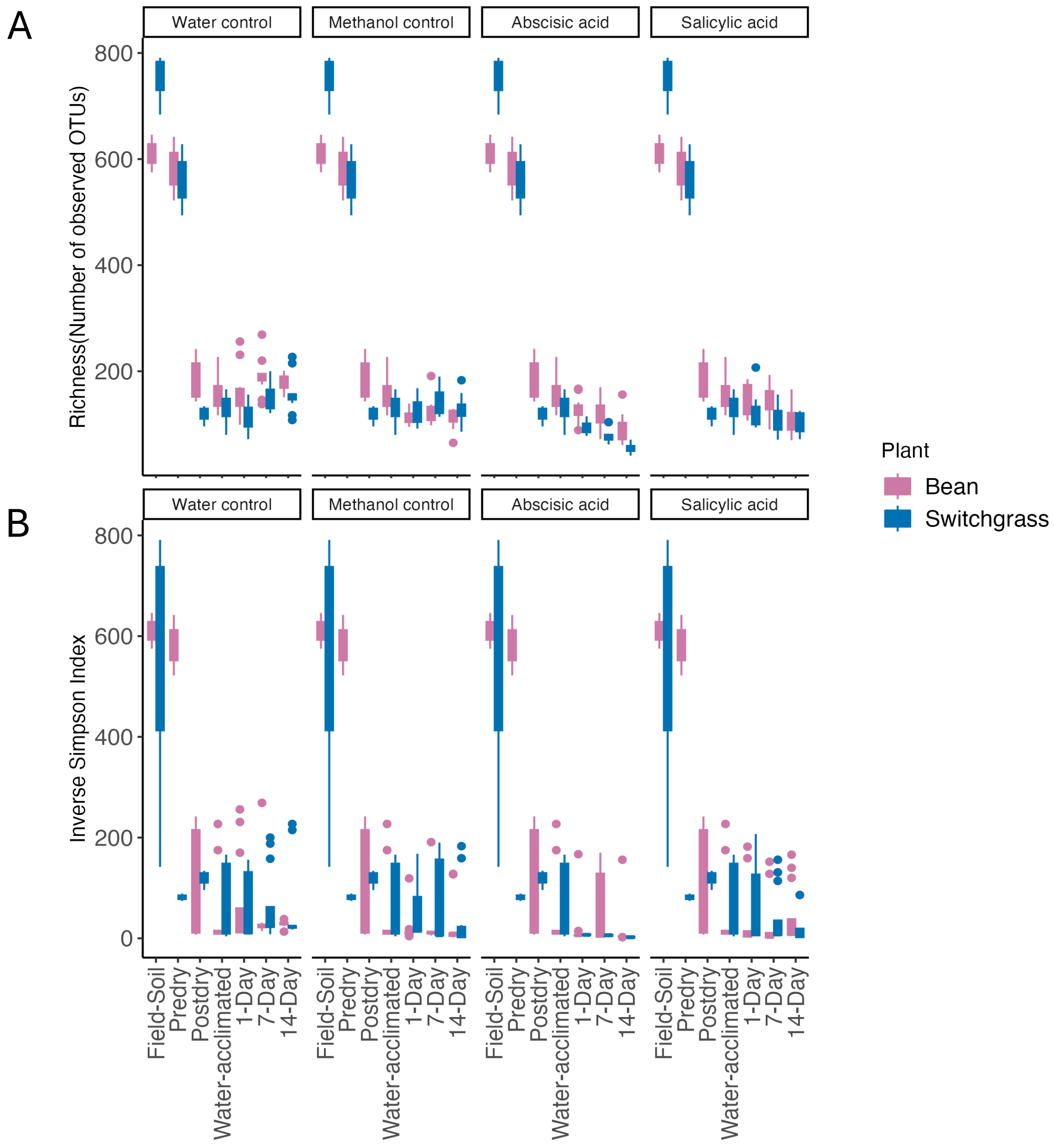


**Figure S6. Changes in active bacterial community richness and diversity over time and as compared to the samples collected before the treatments started**. **A)** Richness decreases with time (starting from water acclimated timepoint) in bean and switchgrass for abscisic acid (ABA) and salicylic acid (SA). **B)** Diversity (Inverse Simpson) decreases with time for switchgrass for ABA and SA.
